# Critical functions and key interactions mediated by the RNase E scaffolding domain in *Pseudomonas aeruginosa*

**DOI:** 10.1101/2024.10.16.618666

**Authors:** Sandra Amandine Marie Geslain, Stéphane Hausmann, Johan Geiser, George Edward Allen, Diego Gonzalez, Martina Valentini

## Abstract

The RNA degradosome is a bacterial multi-protein complex mediating mRNA processing and degradation. In Pseudomonadota, this complex assembles on the C-terminal domain (CTD) of RNase E through short linear motifs (SLiMs) that determine its composition and functionality. In the human pathogen *Pseudomonas aeruginosa*, the RNase E CTD exhibits limited similarity to that of model organisms, impeding our understanding of RNA metabolic processes in this bacterium. Our study systematically maps the interactions mediated by the *P. aeruginosa* RNase E CTD and highlights its critical role in transcript regulation and cellular functions. We identified the SLiMs crucial for membrane attachment, RNA binding and complex clustering, as well as for direct binding to the core components PNPase and RhlB. Transcriptome analyses of RNase E CTD mutants revealed altered expression of genes involved in quorum sensing, type III secretion, and amino acid metabolism. Additionally, we show that the mutants are impaired in cold adaptation, pH response, and virulence in an infection model. Overall, this work establishes the essential role of the RNA degradosome in driving bacterial adaptability and pathogenicity.

**Author summary:** Bacteria must rapidly adapt to changing environments, whether facing temperature shifts, nutrient scarcity, or antibiotic exposure. A key mechanism enabling this adaptability is the regulation of mRNA levels—the molecular blueprints for protein production. This process is governed by the RNA degradosome, a multi-protein complex that processes and degrades RNA to control gene expression. Although the RNA degradosome core function is conserved across bacteria, its composition and organization differ significantly between species, reflecting diverse lifestyles and environmental challenges each bacterium encounter.

In this study, we investigated the RNA degradosome in *Pseudomonas aeruginosa*, a bacterial pathogen causing difficult-to-treat infections in humans. We identified its components and mapped regions within the complex essential for RNA binding, membrane attachment, and spatial organization. Disrupting these regions compromised *P. aeruginosa* ability to survive in cold conditions, respond to stress, and establish an infection. Through this work, we uncovered unique features of the *P. aeruginosa* RNA degradosome that distinguish it from those of other species, emphasizing the RNA degradosome critical role in bacterial adaptability and highlight it as a promising target for therapies against *P. aeruginosa* infections.

## Introduction

In bacteria, the processing and degradation of mRNAs is primarily orchestrated by the RNA degradosome, a multi-enzyme complex that includes ribonucleases, RNA helicases, and various other proteins [1-5]. The RNA degradosome plays a crucial role in enabling bacteria to rapidly adapt to changing environmental conditions by swiftly modifying mRNA levels and broadly regulating gene expression [6, 7].

Most of our knowledge on the RNA degradosome derives from studies in *Escherichia coli*, where the RNase E endonuclease serves as the central component [8-11]. While the catalytic activity of *E. coli* RNase E resides in its N-terminal domain (NTD, amino acids 1-529), the C-terminal domain (CTD, amino acids 530-1061), or scaffolding domain, is intrinsically disordered and contains short linear motifs (SLiMs) of 10 to 40 amino acids to which the other RNA degradosome components bind [11, 12]. Core *E. coli* RNA degradosome components, namely the polyribonucleotide phosphorylase PNPase [13], the RNA helicase RhlB [14], and the glycolytic enzyme enolase [15], interact with RNase E in semi-stoichiometric ratios via the PBS (PNPase binding site), HBS (helicase binding site) and EBS (enolase binding cite) SLiM, respectively [16, 17]. The *E. coli* RNase E CTD also includes an amphipathic helix, called membrane targeting sequence (MTS), which tethers the entire complex to the inner membrane, and at least two RNA-binding SLiMs, AR1 (named also RBD) and AR2, which mediate contact with translating ribosomes, small non-coding RNAs and mRNAs [12, 18-21]. Additional proteins, such as RraA [22, 23], RraB [24], CsdA [25], Ppk [26], MinD [27, 28], and Hfq [29], can interact with *E. coli* RNase E in sub-stochiometric ratios and/or under non-standard conditions of growth (reviewed in [30]). Of importance, the RNase E NTD is essential for *E. coli* growth and rRNA processing [31], whereas deletion of the CTD is not lethal and alters mRNA levels without affecting rRNAs [19, 32-34]. This highlights the dual role of RNase E, with the NTD being crucial for overall cell maintenance, and the CTD regulating specific RNA substrates through the assembly of the RNA degradosome.

RNase E orthologs are present in approximately 49% of sequenced bacterial genomes, and nearly all Pseudomonadota (formerly called Proteobacteria) encode at least one RNase E ortholog, with the notable exception of most *Campylobacter* and *Helicobacter* species [5, 30]. The RNase E NTD is remarkably conserved across most species, while the CTD varies considerably in sequence, length, and SLiMs composition, even between species belonging to the same order [30]. SLiMs variability is thought to drive the diversity of RNase E interacting partners, and therefore RNA degradosome composition, among bacterial species. For instance, in *Caulobacter crescentus,* RNase E interacts with PNPase, RhlB, RNase D, and the Krebs cycle enzyme aconitase [35, 36]. In *Anabaena* sp. strain PCC 7120, RNase E partners with PNPase, enolase, the RNA helicase CrhB, and RNase II [37-39], while in *Pseudomonas syringae* LZ4W, only RNase R and the RNA helicase RhlE were found to co-purify with RNase E in pull-down assays [40]. Similarly, in *Rhodobacter capsulatus*, purification of the RNA degradosome complex via glycerol-gradient centrifugation of cell extracts revealed the presence of RNase E, two unnamed RNA helicases, the Rho transcription termination factor, and two unidentified proteins of 47 and 36 kDa, with only low amounts of PNPase present in the purified fractions [41].

In these examples, the SLiMs mediating protein-protein interactions were not always characterized. Moreover, even when the interaction of RNase E with a certain protein partner is conserved across several species, such as the RNase E-PNPase and RNase E-RhlB interactions, the SLiMs involved in the binding show little sequence identity [30, 36, 37, 42-44]. Therefore, predicting RNA degradosome composition based solely on the RNase E CTD sequence does not seem possible. Experimental characterization of the SLIMs across different bacterial species and their impact on the RNA degradosome composition and function is essential, providing insights into novel regulatory mechanisms that enable bacteria to thrive under stress and adapt to diverse environments.

Little is known about the RNA degradosome composition of the opportunistic human pathogen *Pseudomonas aeruginosa*, which is a leading cause of nosocomial infections worldwide largely due to its remarkable adaptability to different environments and resistance to antibiotics [45-53]. A recent study has revealed that *P. aeruginosa* RNase E co-elutes with PNPase, the PA0428 RNA helicase, and several other proteins in pull-down assays, although their interaction with RNase E has neither been confirmed nor mapped [54]. Our laboratory has characterized the PA0428 RNA helicase, named RhlE2, confirming an RNA-dependent interaction with RNase E and identifying the RhlE2 C-terminal region as necessary and sufficient for this binding [55, 56]. However, the specific RNase E interaction site remained unmapped.

In this study, we characterize the SLiMs within the CTD of *P. aeruginosa* RNase E, identifying those involved in RNA binding (AR1, AR4, and REER-repeats) as well as those mediating interactions with core RNA degradosome components, which we determine to be PNPase and the RNA helicase RhlB (NDPR and AR1, respectively). We also show that RhlE2 interacts with the RNase E NTD in pull-down assays. Protein-protein interactions were probed with multiple experimental approaches, including an analysis of the subcellular localization dynamics of the complex. Additionally, we performed transcriptomic and phenotypic analyses of RNase E CTD mutant strains, revealing the critical role of RNA degradosome complex scaffolding in bacterial virulence, adaptation to cold temperatures and to pH fluctuations.

The ability of *P. aeruginosa* to cause disease is intrinsically linked to a sophisticated gene regulatory network that facilitates bacterial survival and virulence. Within this network, the role of post-transcriptional gene regulation remains only partially explored [57, 58]. By uncovering the composition and functional dynamics of the RNA degradosome in *P. aeruginosa*, our study not only advances the understanding of the molecular mechanisms driving bacterial environmental adaptation but also reveals potential novel drug targets, which could be crucial in the fight against antibiotic-resistant infections.

## Results

### The P*. aeruginosa* RNase E harbours several uncharacterised SLiMs conserved within the Pseudomonadales

*P. aeruginosa* RNase E (Pa RNase E) consists of an NTD (amino acids 1-529) that shares high sequence identity (70%) and similarity (90%) with the NTD of *E. coli* RNase E (Ec RNase E), and a CTD (amino acids 530-1057) that shows only 25% identity and 37% similarity with the CTD of Ec RNase E (Fig. S1). Previous work by Aït-Bara et al. (2015) identified seven different putative SLiMs on the Pa RNase E CTD (MTS, AR4, REE, M29, AR1, M20 and NDPR) [30], which are shown in comparison to the Ec RNase E in Fig. S2A. Of note, the MTS is the only SLiM of Pa RNase E CTD with a propensity to form a secondary structure while in Ec RNase E, the helicase, enolase, and PNPase binding sites also exhibit this feature (Fig. 1 and [12]).

**Figure 1:**
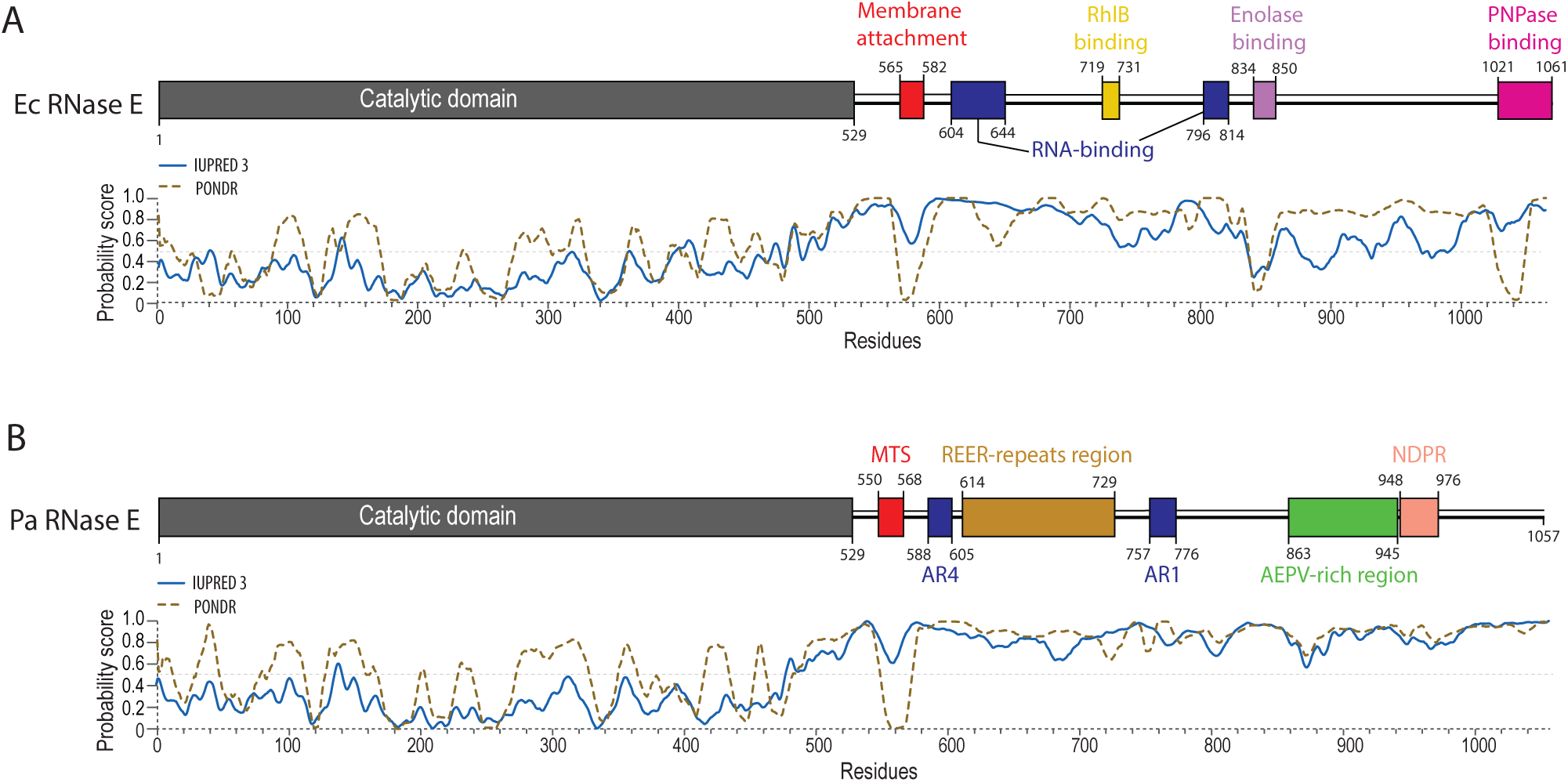
Comparison of *E. coli* (Ec) and *P. aeruginosa* (Pa) RNase E architecture. **(A)** Ec RNase E architecture, adapted from [17]. The NTD is shown is grey, and experimentally defined regions necessary for interaction with the membrane, with RNA, or with the three core protein partners are highlighted in red, dark blue, and yellow/purple/pink, respectively. **(B)** Pa RNase E architecture based on in-silico analyses performed in Fig S1-S2 and in [30]. The SLiM nomenclature was taken from [30], while the consensus sequence characterising each SLiM was specifically adapted based on sequence conservation within the Pseudomonas genus (Fig S1). MTS: membrane-targeting sequence, AR4: arginine-rich motif 4, REER-repeats: region with high frequency of occurrence of “RE[E/D]R” motif, M29: motif 29, AR1: arginine-rich motif 1, M20: motif 20, NDPR: motif with a highly conserved NDPR sequence. All diagrams were drawn to scale. IUPRED3 [95] and PONDR [96] predictions of disorder propensity (probability score) are shown for Ec RNase E **(A)** and Pa RNase E **(B)**.

To refine the prediction of Pa RNase E SLiMs, we performed a multiple sequence alignment of 5000 *Pseudomonas* RNase E orthologs, identifying conserved motifs (Fig. S3). Several highly conserved CTD regions overlap with the SLiMs previously identified, with the notable presence of additional conserved residues located immediately adjacent to these SLiMs (Fig. S2A, Fig. S3). Based on these data, we updated the SLiMs boundaries to include these conserved residues (Fig. 1B, former SLiMs boundaries in Fig. S2A). Upon manual inspection of the CTD sequence, we also observed a large duplicated sequence (residues 635-667 and 682-714) within the CTD region containing the REE and motif 29 SLiMs. Since motif 29 is defined by the conserved motif REERQPR, we merged the two SLiMs and renamed the entire segment (residues 614-729) as the “REER-repeats” region (Fig. 1B). The number of REER repeats varies across *Pseudomonas* species: *P. aeruginosa* strains typically harbour 13 RE[ED]R repeats, whereas the *P. fluorescens*, *P. protegens*, *P. stutzeri*, and *P. syringae* environmental species show a distribution of repeats peaking around 8-10. The exception is *P. putida*, which exhibits a high frequency of 20-23 RE[ED]R repeats (Fig. S2B). This variability suggests a high frequency of indels occurring within the REER-repeats region.

Overall, our in-silico analyses of Pa RNase E CTD sequence refined the number and length of the Pa RNase E SLiMs, highlighting a potential functional significance of the REER-repeats region in *Pseudomonas* species, and suggesting differences in RNA degradosome complex dynamics between *P. aeruginosa* and *E. coli*.

### The REER-repeats, AR1 and AR4 SLiMs mediate RNase E CTD RNA binding and foci formation

Previous works in *E. coli* identified arginine-rich SLiMs as RNA-binding motifs [21, 59]. We therefore hypothesized that the *P. aeruginosa* AR1 and AR4 SLiMs, along with the newly identified REER-repeat region, might be involved in RNA binding. To test this hypothesis, we conducted electrophoretic mobility shift assays (EMSA) to examine the binding of purified RNase E CTD proteins to an *in vitro* synthesized RNA derived from the *malEF* intergenic region (for details, see methods). We tested wild-type (WT) CTD and mutated variants where arginine residues in AR1, AR4, or the REER-repeats were replaced with alanine (named AR1mut, AR4mut, and REERmut, respectively; see Table S4 for a description of the mutations). Qualitative analysis of the EMSA showed that the WT, AR1mut, and AR4mut variants bound RNA with similar efficiency, as indicated by comparable gel shift patterns (Fig. S4A, Fig. 2A). However, the REERmut variant exhibited slightly reduced RNA binding, evidenced by a delayed shift as compared to the former proteins (Fig. S4B and 2A). The combination of AR1 and AR4 mutations resulted in an impaired RNA binding which was further exacerbated when the REER mutation was also introduced: no observable shift in the RNA was detected when incubated with the protein variant carrying mutations in all three SLiMs, even at an RNA-to-protein ratio of 1:80 (Fig. 2A). Together, these results highlight the functional redundancy of the AR1, AR4, and REER-repeat SLiMs in RNA binding, as a complete defect in RNA binding is observed only when all these SLiMs are simultaneously disrupted.

**Figure 2:**
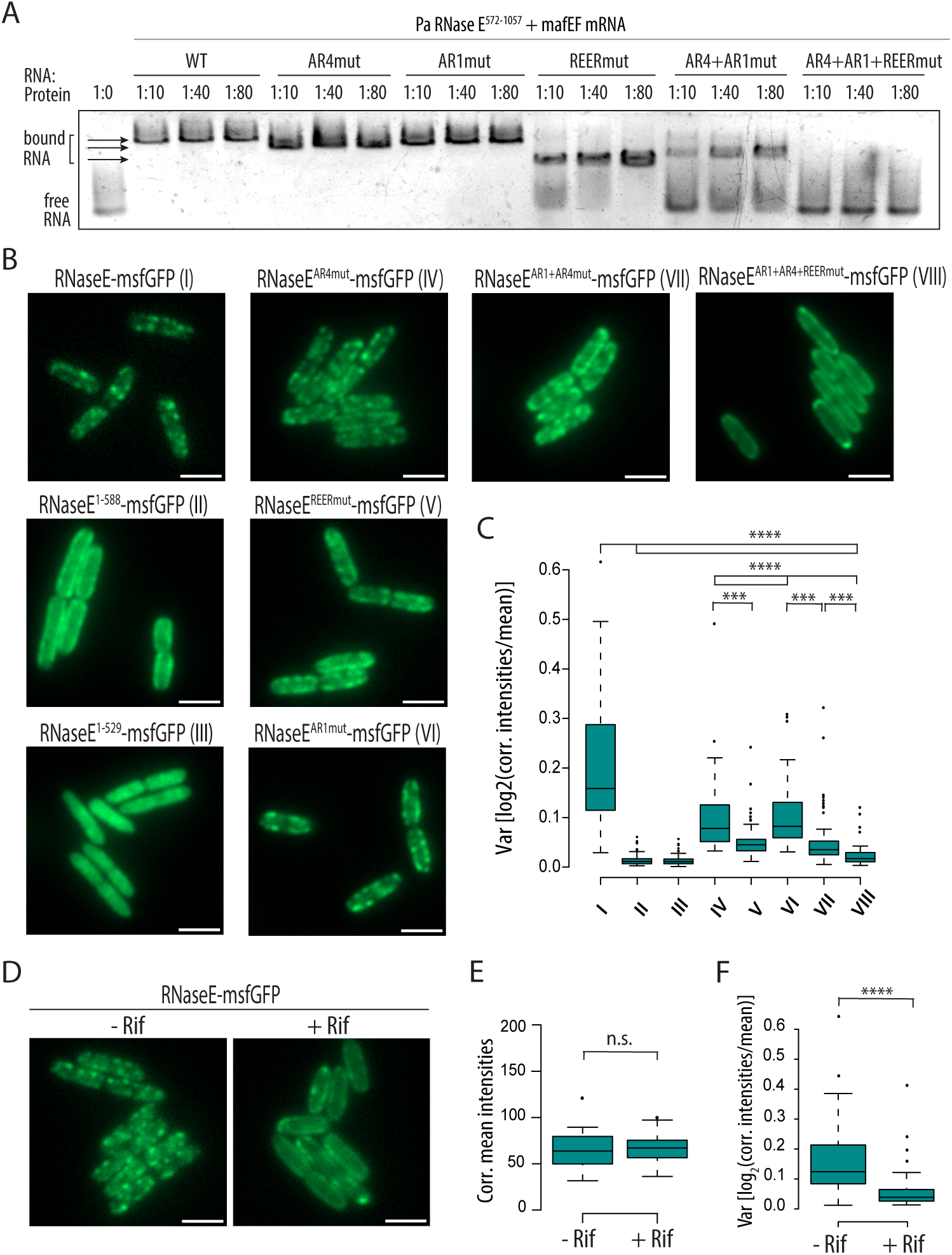
RNA binding of Pa RNase E CTD via the AR1, AR4, and REER SLiMs. **(A)** EMSA with purified His-Smt3 tagged RNase E CTD^572-1057^ variants having either the native sequence (WT) or systematic alanine mutation of the arginine residues found within the REER-repeats region (REERmut), the AR4 SLiM (AR4mut) or the AR1 SLiM (AR1mut). The *malEF* mRNA intergenic region was used as a substrate [97]. Ratios of RNA:protein ranging from 1:10 to 1:80 were tested. See Fig S2 for additional EMSA with additional RNA:protein ratios. **(B)** Representative microscopy images of a PAO1 strain expressing the RNase E-msfGFP fusion or mutated or truncated variants as indicated. **(C)** Quantification of signal intensity variance in the strains of panel B as indication of foci disappearance (see methods). **(D)** Representative microscopy images of a PAO1 strain expressing the RNase E-msfGFP fusion after exposure to rifampicin (+ Rif, exposure to a concentration of 100 μg/mL for 30 minutes) or to DMSO as a negative control (- Rif). Corrected mean fluorescent signal intensity **(E)** and signal intensity variance **(F)** is showed in the graphs (see methods). Each quantification analysis involved at least 50 individual cells. The scale is 2 μm. Statistical significance was assessed by performing t-tests using R. p-values for comparisons tested are indicated as follows: *:<0.05; **:<0.01; ***:<0.001; ****:<0.0001.

One notable characteristic of the RNA degradosome is its tendency to form highly dynamic subcellular foci, also referred to as “clusters,” “puncta,” or “helical structures” [28, 60]. These foci rapidly disassemble upon rifampicin treatment, suggesting their formation is RNA-dependent [60, 61]. In *E. coli*, these foci are membrane-anchored via the MTS SLiM, whereas in *C. crescentus* they remain cytoplasmic, driven by liquid-liquid phase separation [61, 62]. To investigate the subcellular localisation of Pa RNase E, we constructed *P. aeruginosa* PAO1 strains expressing msfGFP-tagged Pa RNase E, Pa RNase E^1-588^, or Pa RNase E^1-529^ from the native *rne* locus. The two RNase E truncations were designed based on our in-silico analysis of Pa RNase E SLiMs, assuming that they would lack the capacity to assemble the RNA degradosome. The key difference between RNase E^1-588^ and RNase E^1-529^ lies in the presence of the MTS (Fig. 1B). Visualization using epifluorescence microscopy revealed that RNase E-msfGFP forms foci concentrated near the cell edge (Fig 2B). Both CTD truncation mutants showed impairment in foci assembly, but while the RNase E^1-588^-msfGFP protein localises uniformly around the edge of the bacteria, the RNase E^1-529^-msfGFP mutant protein additionally fails to localise near the membrane, consistent with the role of the MTS in membrane anchoring (Fig. 2B-C).

Exposure of Pa RNase E-msfGFP expressing cells to high levels of rifampicin (100 µg/ml) led to the dissipation of these foci, resulting in a smoother distribution of the signal along the cell periphery, which phenocopies the localization of the RNase E^1-588^-msfGFP strain (Fig. 2D). Quantitative analysis confirmed a significant reduction in signal intensity variance upon rifampicin treatment as compared to the control (DMSO-only), indicating foci dissipation, while the average signal intensity remained constant, ruling out RNase E-msfGFP proteolysis (Fig. 2E-F, Fig. S4F).

To further verify that Pa RNase E-msfGFP foci formation depends on CTD-mediated RNA binding, we constructed *P. aeruginosa* strains with chromosomally msfGFP-tagged RNase E variants mutated in AR1, AR4, and REER-repeats SLiMs, individually and in combination, as described previously for the EMSA. Visualisation of these variants showed a clear correlation between CTD-mediated RNA binding and foci formation. Specifically, the RNase E AR1+AR4mut-msfGFP showed less distinct foci and lower signal variance compared to RNase E-msfGFP and single AR1 or AR4 mutants (Fig. 2B-C). The RNase E REERmut-msfGFP also had diffuse foci and lower variance, while the RNase E AR1+AR4+REERmut-msfGFP, with abolished RNA binding, displayed the most severe disruption, with nearly complete loss of foci and a very narrow signal intensity distribution (Fig. 2B-C and Fig. S4F). Similar patterns were observed with four other CTD truncations containing different combinations of RNA-binding SLiMs, ruling out the possibility that this foci dissipation is caused by the changes in the charge of the CTD conferred by the Arg to Ala mutations (Fig. S4E).

Of note, we noticed an increased signal intensity in all the msfGFP-tagged RNase E mutants tested as compared to the RNase E-msfGFP (Fig. S4C). Since all constructs are expressed at the native locus, this could likely reflect higher RNase E protein levels, which suggests that CTD-mediated RNA binding might be necessary for RNase E autoregulation [63]. To confirm this, we performed a Western Blot on strains expressing several Strep-tagged RNase E mutants and could indeed observe that RNase E protein levels are increased upon deletion of the CTD or introduction of AR1+AR4+REERmut (Fig. S4D).

Overall, we conclude that in *P. aeruginosa*, the organisation of the RNA degradosome complex into visualisable foci is dependent on the RNase E CTD-mediated RNA binding and suggest that this interaction not only organizes the localisation of the RNA degradosome but could also influence the regulatory activity of RNase E.

### Identification of proteins binding to RNase E scaffold domain

In *P. aeruginosa*, the RNA degradosome was proposed to include the RhlE2 RNA helicase (PA0428), PNPase, and other associated proteins [54, 55]. However, the direct nature of these interactions and the specific interaction sites have not been demonstrated nor mapped. Additionally, RNase R was also suggested to be a *P. aeruginosa* RNA degradosome component, as it is in *P. syringae* LZ4W [30, 40]. To test these hypotheses and identify Pa RNase E CTD interacting proteins, we performed a pull-down assay using *P. aeruginosa* strains expressing C-terminal Strep II-tagged full-length RNase E (RNase E^1-1057^-Strep) or an RNase E truncation missing the entire CTD (RNase E^1-529^-Strep). As an additional control, the assay was also performed with the *P. aeruginosa* PAO1 wild-type (WT) to discard proteins interacting nonspecifically with the Strep-Tactin magnetic beads (Fig. 3A-B).

**Figure 3:**
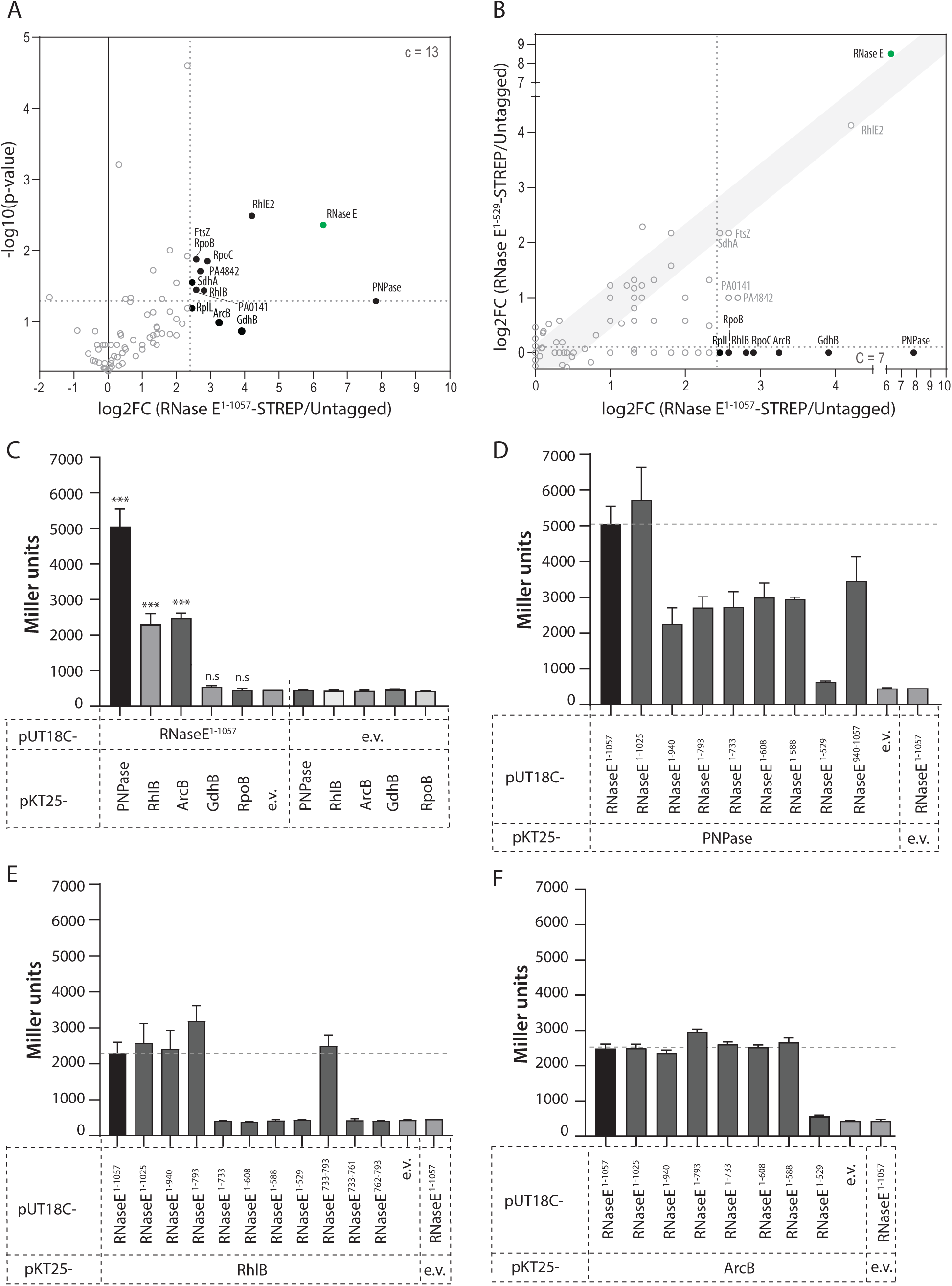
Identification of scaffold domain-dependent RNase E protein partners by pull-down assays and mapping of the interactions by bacterial two-hybrid assays. **(A)** Volcano plot highlighting the relative enrichment of proteins co-eluting with the RNase E^1-1057^-Strep bait protein compared to the untagged negative control. The plot was generated from the average peptide count obtained in duplicate pull-down experiments. An arbitrary threshold of 5.5-fold change was chosen for selection of candidate (c) proteins. **(B)** Scatterplot highlighting comparative enrichment of proteins co-eluting with the RNase E^1-1057^-Strep bait protein (x axis) or with the RNase E^1-529^-Strep bait protein (y axis) compared to the untagged negative control. **(C)** Quantification of β-galactosidase activity in *E. coli* BTH101 cells co-expressing T18-RNase E with T25-PNPase, T25-RhlB, T25-ArcB, T25-GdhB or T25-RpoB. Interaction between T18-RNase E and a T25-tagged protein partner results in a significant restoration of β-galactosidase activity compared with empty vector (e.v.) negative controls. Unpaired Welch’s t-tests was performed using GraphPad Prism to assess differences between every tested protein-protein interaction and the respective negative control. ***: p-value <0.0001; n.s: not significant (99% confidence level). **(D-F)** Mapping of the interaction between RNase E and PNPase **(D)**, RhlB **(E)**, or ArcB **(F)** by quantification of β-galactosidase activity in *E. coli* BTH101 cells co-expressing T25-PNPase **(D)**, T25-RhlB **(E)**, or T25-ArcB **(F)** and either T18-RNase E or one of eight to ten truncated T18-RNase E fragments. Empty vector (e.v.) negative controls are included into each separate graph for comparison.

Eluates of the pull-down experiment were first analysed by silver staining (Fig S5A). The RNase E^1-1057^-Strep eluate revealed a high molecular weight band corresponding to RNase E, along with additional bands absent in the untagged and RNase E^1-529^-Strep samples. These bands likely represent components of the RNA degradosome or cleaved RNase E^1-1057^-Strep intermediates, as RNase E is known to be prone to cleavage [42, 64]. Mass spectrometry (LC-MS/MS) analysis identified a total of 115 proteins present in both duplicates of the RNase E^1-1057^-Strep eluates. 13 proteins (including RNase E) were enriched by at least 5.5-fold as compared with the untagged sample (Fig. 3A). This stringent threshold was chosen since core components of the RNA degradosome should be abundantly associated to RNase E. Of these 12 putative partners, 5 (PNPase, GdhB, ArcB, RhlB and RpoB) were completely absent from both RNase E^1-529^-Strep duplicates and 2, namely RpoC and RplL, were absent from one replicate and present in the second replicate with a peptide count of 2 (Fig. 3B). We selected for further study the first 5 proteins, which were considered the most promising RNA degradosome candidates as their interaction with Pa RNase E seemed fully dependent on the CTD.

It is worth noting that the RhlE2 RNA helicase was, as expected [54, 55], among the proteins enriched in the RNase E^1-1057^-Strep eluate as compared to the untagged control, but surprisingly it was equally enriched in the RNase E^1-529^-Strep eluate condition (Fig 3B), indicating that this interaction occurs via RNase E NTD and not via the CTD.

### RNase E direct interaction with PNPase and RhlB occurs via the NDPR and AR1 SLiMs, respectively

The interaction between RNase E and the 5 candidates was first verified using an adenylate-cyclase-based bacterial two-hybrid (BTH) assay [65]. Co-expression of T25-PNPase and T25-RhlB with T18-RNase E resulted in red colonies on MacConkey agar and a significant increase of β-galactosidase activity compared to empty vector controls (11-fold or 5-fold increase, respectively), confirming their interaction (Fig. S6A, Fig. 3C). Co-expression of T25-ArcB with T18-RNase E also resulted in a significant increase in β-galactosidase activity (5-fold increase) compared to the negative control (Fig. 3C, Fig. S6A). To identify the SLiMs mediating the observed interactions, we generated seven CTD truncations of RNase E, each lacking an additional SLiM as compared with the longer version. Testing these truncations for their capacity to interact with PNPase revealed that only T18-RNase E^1-1025^ showed β-galactosidase activity similar to those of T18-RNase E^1-1057^ upon co-expression with T25-PNPase (Fig. 3D, Fig. S6B). In contrast, all shorter T18-RNase E CTD truncations except T18-RNase E^1-529^, and also T18-RNase E^940-1057^, exhibited a significant increase of β-galactosidase activity as compared with the negative controls but lower than T18-RNase E^1-1057^ (Fig. 3D, Fig. S6B). Overall, this suggests that PNPase interaction with RNase E relies on a complex binding site involving RNase E residues 529-588 and 940-1057.

Co-expression of the T18-RNase E CTD truncations with T25-RhlB revealed that the protein-protein interaction is mediated by residues 733-793: no significant interaction could be seen between T25-RhlB and T18-RNase E^1-733^, although T25-RhlB could interact with T18-RNase E^1-793^ (and every longer CTD truncation) or with T18-RNase E^733-793^ about as strongly as with T18-RNase E^1-1057^ (Fig. 3E, Fig. S6C). Of note, neither T18-RNase E^733-761^ nor T18-RNase E^762-793^ were able to restore β-galactosidase activity above negative control levels when co-expressed with T25-RhlB (Fig. 3E), suggesting that the entire AR1 SLiM, as defined in Fig. 1B, is necessary for interaction with RhlB.

Surprisingly, all T18-RNase E CTD truncations, except T18-RNase E^1-529^, significantly interact with T25-ArcB, suggesting a complex binding that is dependent on residues 529-588 (Fig. 3F, Fig. S6D). Of note, these residues only contain the MTS SLiM, which is presumably involved in membrane binding. Building on the bacterial two-hybrid assays results, we successively employed affinity chromatography with purified proteins to investigate direct binding, minimizing the influence of other protein partners or RNA. Fusion proteins containing a C-terminal 3xFlag tagged RNase E^1-529^, RNase E^572-1057^ (CTD with no MTS) were purified and immobilized on anti-Flag magnetic beads. After extensive washing, the beads were mixed with either purified PNPase or RhlB proteins, and the free or bound fractions were analysed by SDS-PAGE (Fig. 4).

**Figure 4:**
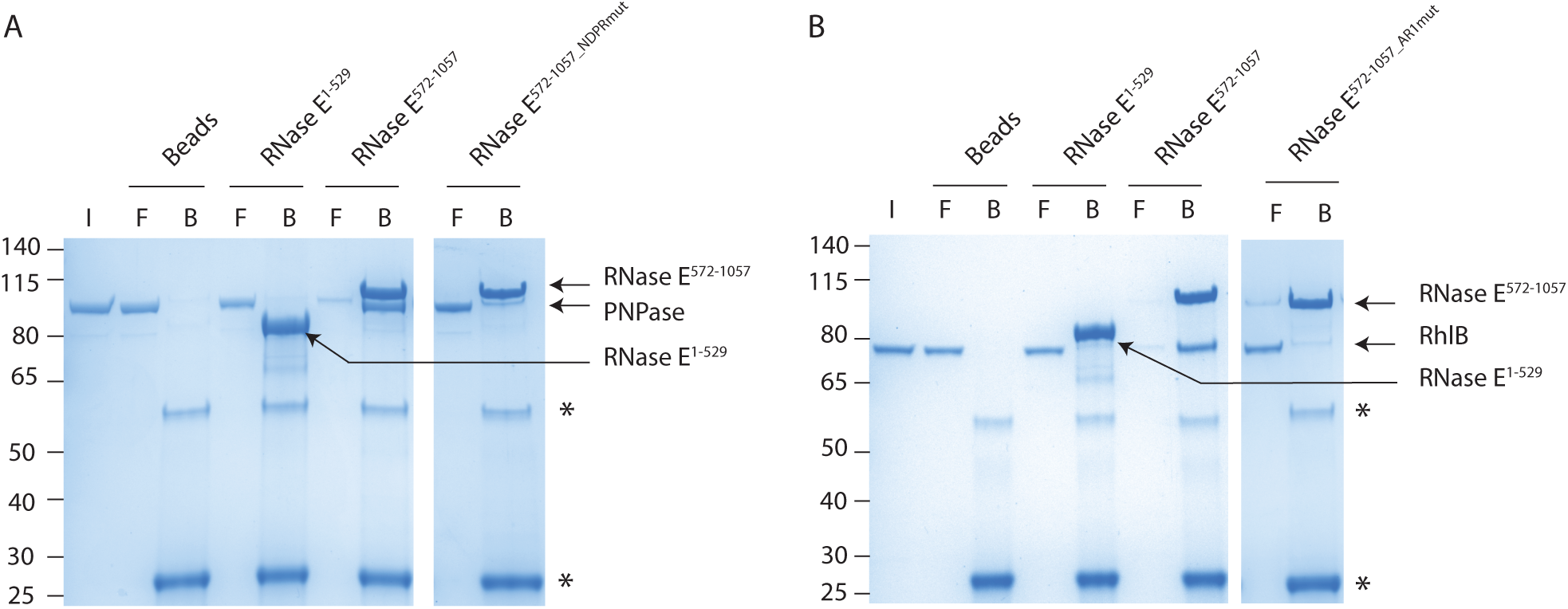
PNPase and RhlB directly interact with RNase E scaffold domain via NDPR and AR1 SLiMs, respectively. Purified Flag-tagged RNase E^1-529^ or RNase E^572-1057^ proteins having either native or mutated sequence were used as baits, immobilized to anti-flag magnetic beads and incubated with purified His-Smt3 tagged PNPase **(A)** or RhlB **(B)** proteins to assess direct protein binding. I: input, F: free fraction, B: bound fraction. NDPRmut, AR1mut: mutations of NDPR and AR1 SLiMs, respectively (see Table S4). Binding assays were repeated at least 2 times and representative gels are shown.

PNPase was retained on beads containing Flag-RNase E^572-1057^, but not on beads alone or containing Flag-RNase E^1-529^ (Fig. 4A), indicating that the RNase E CTD is sufficient for direct interaction with PNPase. Point mutations within the NDPR SLiM (Flag-RNase E^572-1057^_NDPRmut) abolished the binding, confirming that this SLiM is sufficient for direct interaction, as suggested by the BTH assay (Fig. 4A). Of note, PNPase is not retained on beads containing Flag-RNase E^1-588^, which suggests that residues 529-588 are not sufficient to mediate a direct protein-protein interaction, with the NDPR SLiM being the major binding site (Fig. S5B). Similarly, RhlB was retained on Flag-RNase E^572-1057^ but not on Flag-RNase E^1-529^ and point mutations in the AR1 SLiM (Flag-RNase E^572-1057_^AR1mut) showed that this region is crucial for RhlB binding (Fig. 4B).

Contrastingly, ArcB was not retained on beads containing either Flag-RNase E^1-1057^ or Flag-RNase E^1-588^ under the tested conditions, suggesting an indirect type of interaction (Fig. S5C).

To summarize, these experiments clearly establish PNPase and RhlB as core components of *P. aeruginosa* RNA degradosome and demonstrate that the NDPR and AR1 SLiMs are essential for their direct interactions with RNase E.

### PNPase and RhlB are dependent on their physical interaction with RNase E for colocalising in RNA degradosome foci

Given the foci localization of RNase E, we assessed PNPase and RhlB colocalization with RNase E by using *P. aeruginosa* strains carrying *rne::mCherry* and either *pnp::msfGFP* or *rhl::msfGFP* chromosomal fusions. As expected, both PNPase-msfGFP and RhlB-msfGFP formed foci near the membrane and co-localized with RNase E-mCherry in live cells (Fig. 5A). Overlap coefficients between the two fluorescence channels were consistently over 0.9, while Pearson’s coefficients were slightly lower (0.880 for PNPase and 0.773 for RhlB), likely due to differences in signal intensity. Of note, strains expressing any of these chromosomal fusion proteins grow identically to the WT strain at 37°C, suggesting that all the fusion proteins are functional (Figure S7A). We also assessed the subcellular localisation of ArcB-msfGFP and observed a smooth distribution rather than foci, which suggests that the interaction between RNase E and ArcB is likely transient, with ArcB not being a core component of the complex (Fig. S7C).

**Figure 5:**
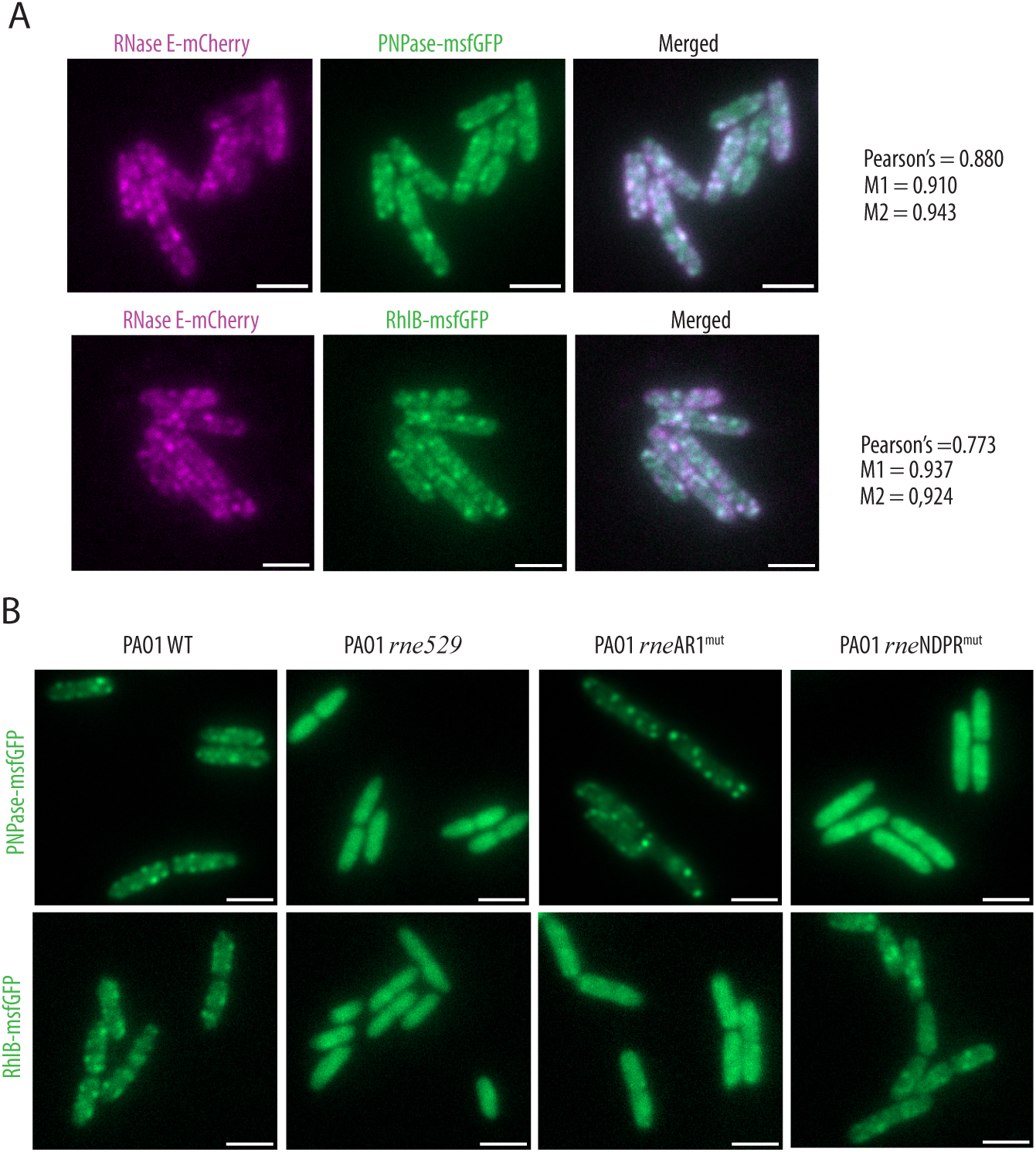
Colocalisation of PNPase or RhlB with RNase E foci is dependent on their interaction with NDPR and AR1 RNase E SLiMs, respectively. **(A)** Visualisation of PNPase-msfGFP (upper panels) or RhlB-msfGFP (lower panels) fusion and RNase E-mCherry fusion co-expressed in the same strain. Pearson’s and M coefficients were calculated in Fiji using the JaCoP plugin [98]. Three independent images were used, each containing at least 30 cells, and an average coefficient was calculated from the three values obtained with each image. **(B)** Visualisation of PNPase-msfGFP (upper panels) and RhlB-msfGFP (lower panels) in strains expressing either the native RNase E protein (full length) or truncated variants, as indicated above each image. Scale is 2 μm.

To further corroborate the need for specific RNase E domains in these interactions, we examined the localization of PNPase-msfGFP and RhlB-msfGFP in strains expressing RNase E^1-1057^ (WT), RNase E^1-529^ (*rne529*), RNase E^AR1mut^ (*rne*AR1^mut^), or RNase E^NDPRmut^ (*rne*NDPR^mut^). Both PNPase-msfGFP and RhlB-msfGFP displayed diffuse cytosolic localization in the *rne529* strain compared to the WT strain (Fig. 5B). In the *rne*NDPR^mut^ strain, PNPase-msfGFP showed a diffuse cytosolic distribution, while RhlB-msfGFP was diffuse in the *rne*AR1^mut^ strain, highlighting an impaired recruitment into the RNase E foci. However, the NDPR or AR1 mutations did not prevent the other protein partner from being recruited into the foci, indicating that both proteins can be independently recruited into RNase E-driven foci (Fig. 5B). We also confirmed that RNase E NDPRmut-msfGFP forms foci, ruling out the possibility that the loss of PNPase foci in the *rne*NDPR^mut^ strain is due to impairment of RNase E-driven foci formation (Figure S7B). These findings confirm the *in vivo* interaction of Pa PNPase and Pa RhlB with the NDPR and AR1 SLiMs, respectively, and demonstrate that these interactions are crucial for their near-membrane foci localization.

### Phenotypic changes associated with partial or complete deletion of RNase E CTD

To investigate the biological importance of RNase E CTD-mediated complex scaffolding in *P. aeruginosa*, we performed some phenotypic assays using the *rne529* and *rne588* strains, as well as strains bearing the AR1 mutation (*rne*AR1^mut^), NDPR mutation (*rne*NDPR^mut^), or AR1+AR4+REER mutation (*rne*AR1+AR4+REER^mut^), abolishing RhlB binding, PNPase binding, or CTD-mediated RNA binding plus RhlB binding, respectively.

On solid media at 37°C, the *rne529*, *rne588* and *rne*AR1+AR4+REER^mut^ strains displayed no severe growth defect but a significantly reduced colony size, suggesting a growth delay (Fig. 6A top panel, Fig. 6B). Indeed, the *rne529, rne588* and *rne*AR1+AR4+REER^mut^ strains exhibited a slight but consistent growth delay at 37°C in liquid culture (Fig. 6C). To verify that the growth delay was caused by truncations/mutations introduced in the *rne* gene, we electroporated the WT or the two most impaired mutant strains with an inducible vector allowing in-trans expression of RNase E, RNase E-msfGFP, or RNase E-mCherry. Cultivation of the strains at 37°C in liquid culture revealed that expression of either native or fluorescently-tagged RNase E could restore identical growth levels compared to the electroporated WT strain, additionally confirming that the fluorescently-tagged RNase E protein is fully functional (Fig S8).

**Figure 6:**
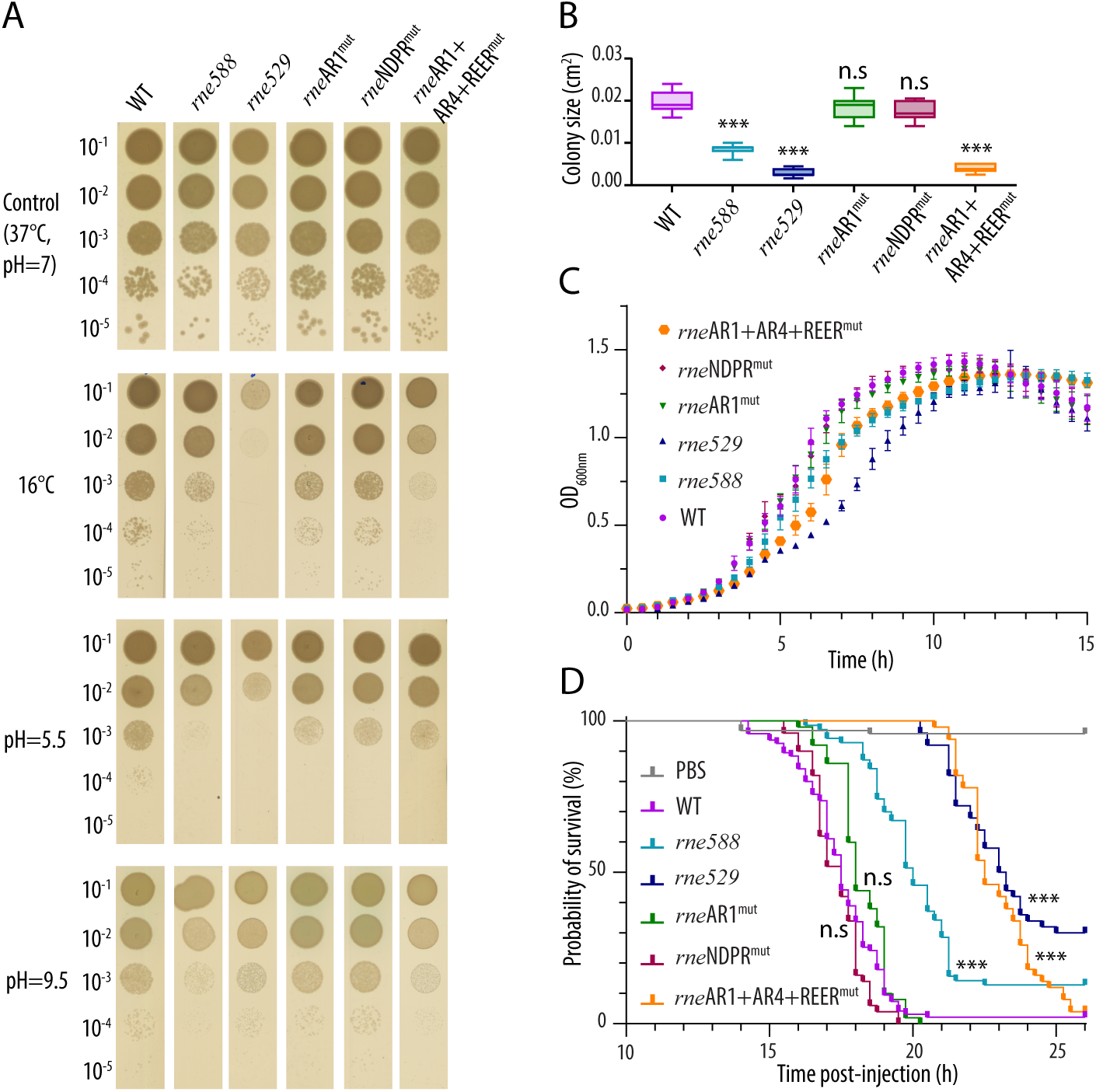
Phenotypes affected by deletion or mutation of RNase E scaffolding domain. **(A)** Growth on NA agar plates at 37°C or 16°C, or under low vs high pH conditions of wild-type (WT), *rne588*, *rne529*, *rne*AR1^mut^, *rne*NDPR^mut^, and *rne*AR1+AR4+REER^mut^ strains, as indicated. Experiments were performed in biological independent triplicates; a representative plate is shown. **(B)** Measured colony size for the different mutant strains as indicated. Quantification was done using the Measure function in Image J after drawing a circle around the edge of each colony. 11 to 13 colonies were quantified for each strain. A one-way Anova test was performed using GraphPad and comparing the colony size of each mutant strain to that of the WT strain. ***: p-value <0.0001; n.s: not significant (99% confidence interval). **(C)** Growth in NYB broth at 37°C in 96-well plates of wild-type (WT), *rne588*, *rne529*, *rne*AR1^mut^, *rne*NDPR^mut^, and *rne*AR1+AR4+REER^mut^ strains, as indicated. Independent biological triplicates, each with four technical replicates, were used to generate the average growth curve shown. **(D)** Killing curves of *G. mellonella* larvae injected with 10 μL of WT or mutant *P. aeruginosa* strains (as in panel A). The control group was injected with 10 μL of sterile PBS instead. Larvae were incubated at 37°C and larval death was monitored over 13-26 hours post-injection. A larva was considered dead when it completely stopped responding to repeated touching stimuli. Two independent replicates were performed using a total of at least 50 larvae for each strain tested. A Kaplan-Meier test for survival analysis was performed using GraphPad Prism and comparing the survival curve of each mutant strain to that of the WT strain. ***: p-value <0.0001; n.s: not significant (99% confidence interval).

Interestingly, when cultured on solid media at 16°C, the *rne588* strain showed impaired growth, while the growth defects of the *rne529* and *rne*AR1+AR4+REER^mut^ strains became more pronounced, with *rne529* barely growing (Fig. 6A, upper middle panel). Since the *rne*AR1^mut^ and the *rne*NDPR^mut^ strains exhibited no significant differences compared to the WT, these results indicate that the cold sensitivity of the *rne588* strain can be attributed to the loss of RNase E CTD RNA binding. Moreover, the severe growth defect observed in the *rne529* mutant indicates that additional delocalisation of RNase E from the membrane exacerbates the cold sensitivity caused by deletion of RNase E RNA-binding SLiMs.

Besides low temperature, we also tested sensitivity to low or high pH (Fig. 6A, lower panels): while growth at pH≃9.5 is rather similar between the different strains, deletion of the CTD (*rne529* and *rne588*) impairs growth at pH≃5.5, although the three CTD point mutant strains are mostly unaffected. This suggests that, although loss of interaction with PNPase or RhlB separately does not seem to impair growth, the combined loss of RNase E protein-protein and protein-RNA interactions results in a growth defect under low pH conditions. Finally, we assessed whether the RNase E CTD mutants might be impaired in virulence using *Galleria mellonella* larvae, a highly predictive infection model for studying mammalian infection processes [66]. Infection with PAO1 WT, *rne*AR1^mut^ and *rne*NDPR^mut^ strains resulted in nearly 100% larval mortality within 20 hours post-injection (Fig 6D). In contrast, the larvae infected with the *rne588* mutant exhibited 50% survival at this time point, while those infected with *rne*AR1+AR4+REER^mut^ or *rne529* mutants showed 100% survival, with larval death occurring only after 21 hours post-injection (Fig 6D). These results strongly indicate that the ability of *P. aeruginosa* to infect a host is compromised when RNase E CTD-mediated RNA binding is disrupted. This impairment is likely attributable to several contributing elements: reduced bacterial growth within the host, deficiencies in virulence factor production, and a decreased ability to endure host immune responses and host-related stresses.

### Effect of RNase E CTD on *P. aeruginosa* transcripts steady-state levels

To better understand the phenotypes observed and evaluate the impact of RNase E CTD loss on *P. aeruginosa* transcriptome, we conducted RNA-sequencing analysis on *P. aeruginosa* WT, *rne529,* and *rne588* mutant strains grown at 37°C until the late-exponential growth phase (OD_600nm_ of 1-1.2). Biological triplicates were analysed and show a high degree of relatedness within each strain (Fig. S9A-B). Under these conditions, the loss of the entire RNase E CTD in the *rne529* strain affected the steady-state levels of 450 transcripts, with 183 downregulated and 267 upregulated (adjusted p-value < 0.05, fold change |FC| ≥ 2) (Fig. 7A). In the *rne588* mutant, 417 transcripts were differentially expressed, with 194 downregulated and 223 upregulated (adjusted p-value < 0.05, fold change |FC| ≥ 2) (Fig. 7B). Differential expression analysis on reads mapping to predicted small RNA genes (sRNAs) identified 14 predicted sRNA genes with significantly enriched reads in the *rne529* mutant and 5 in the *rne588* strain (Table S5).

**Figure 7:**
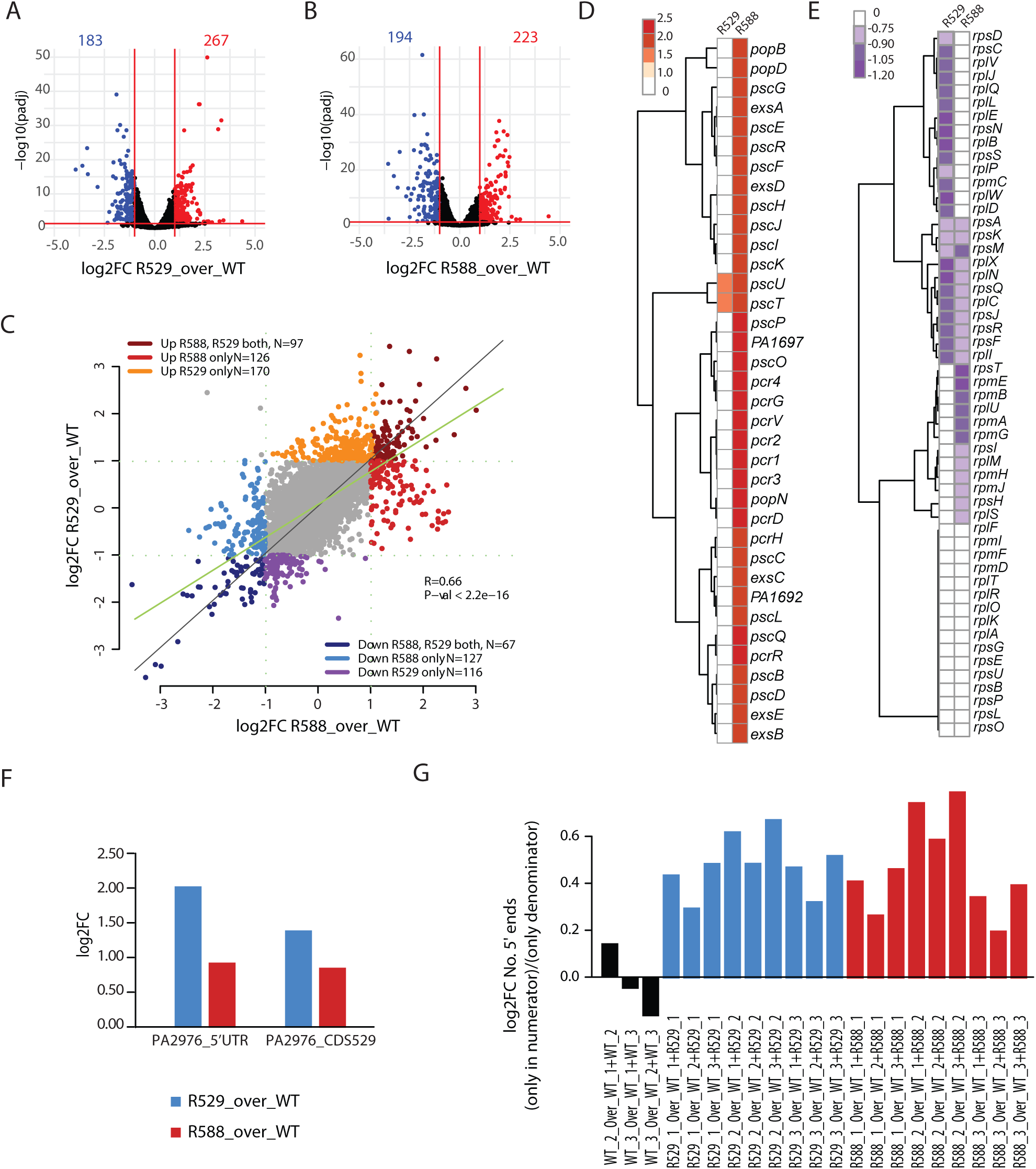
RNA-seq analysis of RNase E CTD truncated mutants reveals broad transcriptome changes. (A-B) Volcano plots highlighting genes differentially regulated in the **(A)** *rne*^1-529^ (R529) or **(B)** *rne* ^1-588^ (R588) mutant strains compared with the WT strain. Genes significantly up- or down-regulated (adjusted p-value < 0.05, fold change |FC| ≥ 2) are highlighted in red or blue, respectively. **(C)** Scatterplot of genes differentially regulated in *rne* ^1-529^ strain versus genes differentially regulated in *rne*^1-588^ strain, highlighting commonalities and differences between the two mutant strains. **(D-E)** Clustered heat maps of type 3 secretion system **(D)** or ribosomal proteins **(E)** transcript levels in the RNase E^1-529^ and by about 2-fold in the RNase E^1-588^ strains as compared to the WT strain (fold change). **(F)** Differential analyses (log 2-fold change) of normalized total reads counts corresponding to the 5’ UTR region of the PA2976 (*rne*) transcript or RNase E NTD coding sequence in the RNase E^1-529^ and by about 2-fold in the RNase E^1-588^ strains as compared to the WT strain. **(G)** Proportion of loci with 5’ read ends (sequenced fragments) counted only in a strain (numerator) relative to the 5’ read ends counted in that strain and in a WT replica (denominator). A total of 8 million reads was analysed in each strain to exclude an effect due to the library depth (see methods). Histograms of individual values calculated per each strain and replicate is shown in Figure S10.

A comparison of the *rne529* and *rne588* strains revealed that 164 mRNAs were consistently regulated in both mutants (97 upregulated and 67 downregulated, see Figure 7C). Among the strongest commonly downregulated genes were: (i) the *pqsABCD*, *pqsE*, and *pqsH* genes, involved in the synthesis of quorum-sensing signal molecules alkylquinolones PQS (2-heptyl-3-hydroxy-4(1H)-quinolone) and HHQ (2-heptyl-4(1H)-quinolone), which play roles in virulence [67], (ii) the *phzA1B1* and *phzA2B2* phenazine biosynthesis genes, which are important for synthesis of the toxic secondary metabolites phenazine and pyocyanin [68], and (iii) the *agtABCD* gene cluster for 4-aminobutyrate (GABA) and 5-aminovalerate (AMV) uptake [69]. Surprisingly, the two strains exhibited more differences in mRNA levels than shared changes. In the *rne588* strain, the mRNA levels corresponding to type 3 secretion system genes, which deliver toxins to host cells [70], were consistently upregulated, while they did not vary significantly in the *rne529* strain as compared to the WT (Fig 7D). On the other hand, ribosomal and many amino acid metabolism genes (tryptophan, arginine, alanine, aspartate and glutamate) were downregulated only in the *rne529* strain, which could correlate with the general growth defect observed (Fig. 7E and Table S5).

Considering the autoregulation of RNase E reported in *E. coli* and *C. crescentus* [63, 71], we quantified reads mapping to the 5’ untranslated region (UTR) of the *rne* gene (transcription start site determined in [72]) and observed increases of 2 and 0.96 log_2_ folds in the *rne529* and *rne588* mutants, respectively, compared to the WT (Fig. 7F). Similarly, we observed a 1.43 and 0.88 log_2_ fold increase in the transcript corresponding to the NTD of RNase E in the *rne529* and *rne588* mutants, respectively (Fig. 7F), in agreement with the observed increase in protein levels (Fig. S4C-D).

Finally, to assess RNA degradation efficiency in our strains, we analysed the 5’ reads ends of sequenced RNA fragments. As previously performed in other studies [56, 73], we calculated the ratio between the number of loci with 5’ ends present only in a mutant and not in the WT, over the sum of loci with 5’ read ends seen in both mutant and WT. We performed the analysis in the three genetic backgrounds (WT, *rne529*, *rne588*) and for each replica independently to ensure consistency (see methods and Fig S10). Despite the background of random fragmentation intrinsic to the sequencing protocol, a consistent increase on the number of 5’ ends in the *rne529* and *rne588* strains was observed as compared to the WT (Fig. 7G), suggesting an accumulation of RNA degradation intermediates and an altered RNA degradation in the RNase E truncated mutants.

## Discussion

The composition and the functions of the RNA degradosome vary among bacterial species, likely reflecting diverse lifestyles and niche adaptations [30]. Studying the RNA degradosome in various bacterial species is therefore crucial for elucidating how bacteria respond and adapt to different environmental conditions and sustain various types of stress. In this study, we characterized the RNA degradosome of *P. aeruginosa*, a versatile opportunistic pathogen and widely used model organism for bacterial pathogenesis and environmental adaptation.

In the first part of our study, we focused on the putative SLiMs within the CTD of *P. aeruginosa* RNase E and assessed their contribution to RNA binding and foci assembly. We successively identified RhlB and PNPase as core RNA degradosome components, as well as the SLiMs mediating these interactions (Fig. 8). A notable feature of SLiMs is their poor sequence conservation, even when they mediate interactions with the same protein partner. Our findings confirm this variability for both PNPase and RhlB interactions.

**Figure 8:**
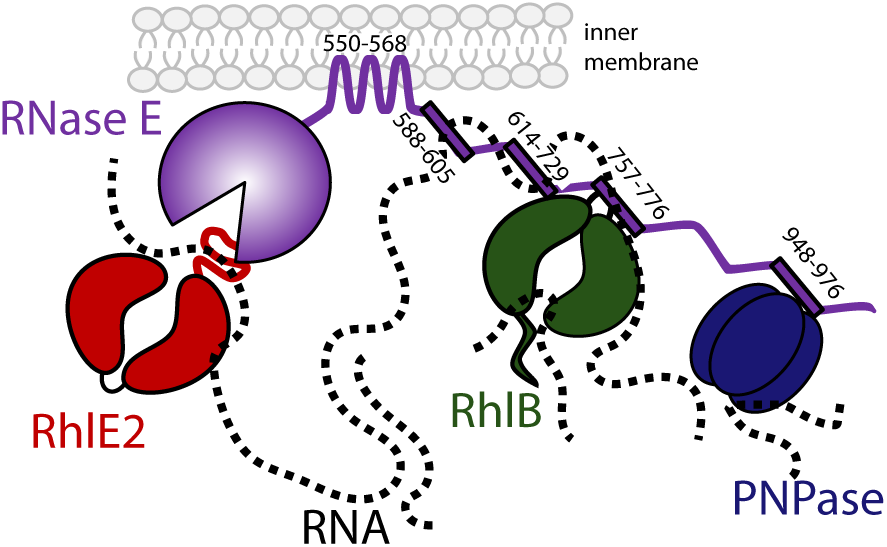
Model of RNA degradosome in *P. aeruginosa*. Numbers corresponds to RNase E SLiM start-end residues involved in membrane, RNA, RhlB or PNPase interaction. RNA is represented as a dotted line. See discussion for details.

PNPase, a known partner of RNase E in several species, binds through SLiMs with significantly different sequences across organisms. For instance, in *Anabaena* PCC7120, the interaction occurs via a cyanobacterial SLiM having the conserved residues RRRRRRSSA [37], while in *E. coli* or *C. crescentus*, the sequences are WQRPTFAFEGKGAAGGHTATHHASA [30] and APPEKPRRGWWRR [43], respectively. Our pull-down assays and *in vitro* binding assays showed that PNPase interacts with RNase E in *P. aeruginosa* via the NDPR SLiM, which has yet another distinct sequence (TGRALNDPREKRRLQREAERLAREAAAAA). These sequence variations may reflect species-specific interactions, as suggested by experimental work showing lack of heterologous interactions between RNase E and PNPase of *E. coli* and *Pseudoalteromonas haloplanktis* [42]. Despite sequence variability, the PNPase interacting SLiM is consistently located towards the end of the RNase E CTD [30, 37, 43], suggesting convergent evolution in the positioning of the PNPase relative to other RNA degradosome components and to RNase E catalytic domain.

Of note, a recent study on a *P. aeruginosa* mutant with a 50 bp deletion in the 3’ end of the *rne* gene revealed a hypervirulence phenotype, characterized by enhanced cytotoxicity and siderophore production [74]. Based on *E. coli* RNA degradosome model, this phenotype was associated with a possible impairment of PNPase binding to Pa RNase E or a different RNA degradosome activity [74]. Our data reveal that PNPase can interact with the RNase E^1-1025^ truncation (BTH assay), ruling out the former hypothesis. However, we cannot exclude the possibility that this region might be involved in interaction with a yet unknown partner.

Regarding RhlB, we show that in *P. aeruginosa*, this RNA helicase directly interacts with RNase E via the AR1 SLiM (RPRRRSRGQRRRSNRRERQR). Mutation of AR1 disrupts not only the RhlB-RNase E binding *in vitro*, but also RhlB subcellular localisation near the membrane and co-localization with RNase E *in vivo*. Interestingly, in *E. coli* and other species the AR1 SLiM mediates RNA interaction [30, 42]. In *P. aeruginosa*, AR1 also contributes to RNA-binding in addition to RhlB binding. Given that in *E. coli,* a single arginine mutation within the HBS SLiM (R730A) is sufficient to disrupt RhlB binding [75], it is possible that only one or a few arginine residues of the AR1 SLiM are involved in interaction with RhlB, while the others mediate RNA binding. Further experiments are needed to address this hypothesis and to understand the dynamics of the RhlB-RNA-RNase E complex in *P. aeruginosa*. Moreover, it has been shown that the RNA-dependent ATPase and unwinding activity of RhlB are stimulated by binding to RNase E [76]. Future studies will investigate whether it is also the case in *P. aeruginosa*.

In contrast with the *E. coli* RNA degradosome, we did not find any evidence of an interaction between RNase E and enolase in *P. aeruginosa* when the bacteria grow in rich medium and standard laboratory conditions. Although our pull-down and bacterial-two hybrid data indicate a probable interaction between RNase E and ArcB, a catabolic ornithine carbamoyltransferase from the arginine deaminase pathway [77], no direct protein-protein binding could be observed *in vitro,* nor were the two proteins colocalising in vivo. We propose that RNase E indirectly interacts with ArcB, probably in a RNA-dependent manner. Indeed, the *P. aeruginosa arcDABC* polycistronic transcript undergoes endonucleolytic cleavage at multiple sites for its processing [78], suggesting that an regulatory interaction between RNase E and ArcB polypeptide could occur during co-translational RNA cleavage [18].

Additionally, the RNase E-based RNA degradosome of *P. haloplanktis, P. syringae*, and *R. capsulatus* do not seem to contain either a metabolic enzyme upon standard growing conditions [40-42]. This lack of RNase E interaction with metabolic enzymes may reflect distinct physiological roles of the RNA degradosome in different bacterial species, potentially linked to specific metabolic adaptations or regulatory mechanisms that modulate RNA stability and processing depending on nutrient availability. However, we cannot exclude that some protein partners were missed as we chose a stringent threshold for analysis of the pull-down results. Transient or condition-specific accessory protein partners remain to be identified, and compositional biases such as the conserved AEPV-rich part of the RNase E CTD are not yet understood, as no ligand or biological function has been assigned to this region even in well-studied RNase E orthologs.

It is also important to mention that throughout this study, we chose to focus on core RNA degradosome components that are identified based on their binding to RNase E CTD, but we cannot exclude that some functional interactions occur with the NTD. For example, RhlB was found to interact with RNase E NTD in *C. crescentus* [36], while in *Anabaena* the RNA helicase CrhB, RNase II and enolase were shown to associate with RNase E via its NTD [38, 39]. Similarly, we found that the *P. aeruginosa* RhlE2 RNA helicase, previously shown to associate with RNase E in an RNA-dependent manner [55], interacts with the N-terminal domain of Pa RNase E. Although RhlE2 is not considered a core RNA degradosome component based on our assumption of CTD-mediated interactions, this association remains functionally significant [55, 56].

In the second part of our study, we evaluated the importance of RNA degradosome assembly at both transcriptomic and phenotypic levels through an analysis of *P. aeruginosa* strains carrying RNase E CTD truncations and mutations. Comparing the *rne588* and *rne529* strains allows us to discuss the importance of RNA degradosome assembly and membrane attachment in *P. aeruginosa*. The two mutants exhibited varying degrees of sensitivity to cold growth, with *rne529* being the most affected, indicating that cold growth requires both RNase E membrane attachment and RNA degradosome assembly. Loss of the CTD without delocalisation from the membrane (*rne588*) does not affect growth in *P. aeruginosa* under standard laboratory condition (i.e., 37°C and rich medium), but impacts its capacity to sustain growth in the cold. Transcripts levels were also generally affected in the mutants, as well as global RNA degradation, as shown in the 5’ read ends analysis. Decreased degradation of cleaved transcripts could lead to an upregulation of their levels. Therefore, direct and indirect effects remain to be distinguished and those changes remain to be correlated with proteins levels.

Surprisingly, we did not find any phenotypic changes associated to the loss of RhlB or PNPase binding, questioning the importance of the RNA degradosome protein association for the complex functionality. Competition assays could be useful in identifying any fitness defects in our mutants, as done in *E. coli* for RNase E CTD truncations [19, 62], or different phenotypes or stresses could be tested.

On the other hand, our phenotypic analysis highlights the crucial role of CTD RNA binding for the functionality of the RNA degradosome complex. Mutations in the SLiMs responsible for RNA binding (AR1, AR4, and the REER-repeats) not only impair *in vivo* RNA degradosome foci assembly and/or stability but also lead to decreased *P. aeruginosa* virulence and reduced growth at both 37°C and cold temperatures. The effect of these mutations on foci formation mirrors the effect observed upon cell exposure to rifampicin, which decreases RNA levels in the cell, and aligns with previous studies showing that Ec and Cc RNase E orthologs also assemble foci in a CTD-dependent and RNA-dependent manner [60-62]. Our data thus suggest that impairing foci formation has a major effect on the activity of the RNA degradosome in *P. aeruginosa*.

Overall, our study identified core RNA degradosome components in *P. aeruginosa* and underscores the role of this complex in bacterial adaptation to environmental changes and pathogenesis. By revealing *P. aeruginosa*-specific interactions and their functional significance, our findings provide valuable insights into the molecular mechanisms of adaptation in this pathogen and highlight the potential of the RNA degradosome as a drug target.

## Methods

### Bacterial strains and growing conditions

A list of bacterial strains used can be found in Table S1. Unless otherwise specified, *P. aeruginosa* PAO1 was grown at 37°C in NYB (25 g of Difco nutrient broth, 5g of Difco yeast agar per liter) with shaking at 180 rpm or on Nutrient Agar (NA) (40 g Oxoid blood agar base, 5 g Difco yeast agar per liter). When required, antibiotics were added to these media at the following concentrations: 100 µg/ml ampicillin, 50 μg/mL kanamycin, 20 μg/mL tetracycline and 10 µg/ml gentamicin for *E. coli*; and 50 μg/mL or 100 μg/mL tetracycline and 50 µg/ml gentamicin for *P. aeruginosa*.

### Construction of plasmids used in this study

Information about plasmid construction and primers used in this study can be found in supplementary tables S2 and S3, respectively. Plasmids pUT18C and pKT25 were generally cloned by restriction enzyme digestion using EcoRI-HF and BamHI-HF (NEB). When a gene to be inserted could be internally cleaved by EcoRI and/or BamHI, the enzymes MfeI and/or BglII, respectively, were chosen instead during primer design to allow ligation of the insert to the vector digested with EcoRI/BamHI thanks to compatible cohesive ends. For cloning of *gdhB* into pKT25, NEB assembly^®^ was used instead, due to the presence of many restriction enzymes internal cleavage sites. Ligation reactions were carried our using T4 DNA ligase (Roche or NEB) at 16°C overnight, or at room temperature for 2-3 hours, following standard protocols. When cloning inserts into the suicide vectors pME3087 or pEXG2, the NEBuilder HiFi DNA assembly Master Mix (NEB) was used according to the manufacturer’s recommendations. To constructs RNase E CTD mutants, the DNA was synthesized by GeneArt (ThermoFisher Scientific) and cloned into appropriate vectors for protein purification or strain construction. A description of the introduced mutations can be found in Table S4. To construct pME6032-based expression vectors, restriction enzyme digestion was carried out using EcoRI and KpnI for both vector and inserts.

Ligation mixtures or the assembly reactions were transformed into chemically competent *E. coli* DH5α cells by heat-shock, plating on appropriate selective LB agar medium. The resulting colonies were screened by PCR for the presence of the insert in the plasmid of interest. After plasmid extraction (Miniprep kit, Sigma-Aldrich/Merck), plasmids were verified by Sanger sequencing at Microsynth.

### Construction of *P. aeruginosa* mutant strains

To construct chromosomal gene truncations or insert a Strep-tag or fluorescent protein into the chromosome, in frame with a gene of interest, the suicide vectors pEXG2 [79] or pME3087 [80] were used. Briefly, approximately 500 bp regions located immediately upstream or downstream of the region to be edited, and, when applicable, the tag or fluorescent protein of interest, were amplified by PCR and inserted together into the vector backbone using NEB assembly^®^. After sequencing verification of the constructed plasmid quality, the plasmids were introduced into *P. aeruginosa* PAO1 strain by triparental mating using the *E. coli* strain HB101 (pRK2013). After selection with the appropriate antibiotic (tetracycline for pME3087 or gentamicin for pEXG2), merodiploids were resolved as described in [81] for pME3087 or [79] for pEXG2, and strains with the desired insertion/deletion were identified by colony PCR.

### RNase E sequence analysis

Group-specific homologs of RNase E in *P. aeruginosa*, *P. fluorescens*, *P. protegens*, *P. putida* and *P. stutzeri* were retrieved from NCBI databases using blastp [82] with *P. aeruginosa* PAO1 RNase E as a query; only sequences with query coverage > 0.5 were conserved (total of 5000 proteins). The number of RE[E/D]R motifs in each protein sequence was determined using the vcountPattern function from the “biostrings” R package [83]. The similarity of the protein sequences within a taxonomic group was determined using the conserv function (method=“similarity”) from the bio3D R package on a muscle alignment of the protein sequences[84, 85].

### Electrophoretic motility shift assay

The *malEF* transcript was synthesized with a T7 transcription kit (Invitrogen) on linear DNA template (172 bp) obtained by PCR with primers p253-p254 (see Table S3). The electrophoretic mobility shift assay (EMSA) was performed following the protocol of [86]. Briefly, the reaction mixtures (15 μl) contained 35 nM of RNA, 50mM Tris-HCl, 100mM NaCl, 5mM DTT, 1.0 units/μl RNasin (Promega), and His10Smt3-RNase E proteins as specified, with protein concentration expressed as a molar ratio of relative to RNA concentration (see Fig. 2A and S4A-B).

The reactions were incubated at 4 °C for 30 min before loading on 2% agarose gel and run at 150 V for 15 min. RNA bands were visualized by gel staining with SybrGold (Invitrogen).

### Epifluorescence microscopy imaging

Phase contrast and fluorescence microscopy were performed on a Zeiss Axio Imager 2 equipped with an objective alpha Plan-Apochromat 100x/1,46 Oil Ph3 M27 and a ZEISS Axiocam 305 mono CCD camera. Bacteria were grown until OD_600nm_= 0.2-0.6 (approx. 2.5 hours) in 3 ml of NYB and were then placed on an agarose pad containing 1% agarose dissolved in 0.5X PBS. Images were processed with Fiji (NIH, USA) [87].

### Image analysis

All image treatments and analysis were performed using Fiji (NIH, USA)[87]. Representative images shown had contrast adjusted for display by opening the images in Fiji and using the Reset function in the Adjust Brightness/Contrast menu, after average background intensity subtraction using the Math -> Subtract function. These average background intensity values were obtained by measuring a square area devoid of any cells.

For quantification of signal intensity in Fiji, a line was drawn along the longitudinal axis of bacterial cells and signal was measured using Plot Profile. Results of this function were saved as .csv files. For each strain, at least 50 bacterial cells were measured from the raw images, using at least two different images for quantification. Average background values were subsequently subtracted from each results file using a .csv file containing the Plot Profile measurements of a linear area devoid of cells for each image used. Boxplots of fluorescence intensity and signal variance or pooled histograms were plotted using R [88]. The signal variance was obtained by calculating the variance of the log_2_ fold-change between each measured value and the average of all values for a given measured cell. This was done to normalize the data and obtain comparable variances without introducing a bias due to differences in overall signal intensity.

### Pull-down identification of RNase E partners

Fresh cultures of PAO1 WT, PAO1 RNase E_1-529_-Strep, or PAO1 RNase E_1-1057_-Strep were inoculated to OD=0.05 from overnight cultures in 500 mL Erlenmeyer flasks. Cultures were grown until approx. OD_600nm_=0.8-0.9. 50 mL of each culture was harvested by centrifugation at 4500 *x g* for 10 minutes. Pellets were stored at -80°C until needed.

Each thawed pellet was then resuspended in 3,25 mL of lysis buffer (for 10 mL of buffer, prepared just before use: 20 mM Tris (pH 10), 250 mM NaCl, 10% v/v glycerol, 1 EDTA-free protease inhibitor tablet, 50 μg/mL lysozyme). Samples were incubated the samples on ice for about 30 minutes, transferred into 15-mL Falcon tubes and sonicated using a thin probe for 6x30s, with 15 seconds break in between. A pulse of 70% and a power output of 50% were used. After sonication, 32.5 μL of Triton X100 10% were added to each sample (∼0.1% final) before incubation on a rotary wheel at 4°C for 8 minutes. The samples were transferred into 2 mL Eppendorf tubes and centrifuged for 1 hour at 10500 *x g* at 4°C. MagStrep^®^ Strep-Tactin^®^XT beads (IBA, cat# 2-5090) were washed in wash buffer (20 mM Tris (pH 10), 250 mM NaCl, 10% v/v glycerol) according to the manufacturer’s instructions. One mix of beads was prepared for all the samples with a volume of beads suspension of 45 μL per sample. Final resuspension was done in 200 μL wash buffer per sample, plus an additional 10 μL. After centrifugation of the samples, the supernatant of each sample was regrouped into a 5 mL Eppendorf tube, and 200 μL of washed beads suspension were added per sample. Incubation on a rotary wheel was done for 2h15 at 4°C. The 5 mL tubes were then placed against a magnetic rack and all the supernatant was carefully removed. A first wash volume of 500 μL was added gently and agitated by horizontal rotations of the tubes. Tubes were placed back against the magnetic rack and the wash volume was removed. A second wash step was performed identically with a volume of 225 μL instead of 500 μL. A third wash step of 225 μL was also performed, and the resuspended beads were transferred into a clean 1.5 mL Eppendorf tube by gentle pipetting with a 1000 μL tip, before removing the wash volume. The beads were covered with 10 μL of sterile PBS and stored at 4°C before LC-MS/MS analysis at the Proteomics Core Facility (University of Geneva). For plots in Fig. 3A-B, the log2 fold-change (log2FC) was calculated using the following formula: log2(average peptide count of protein *x* identified in the RNase E^1-xxx^-Strep / average peptide count of protein *x* identified in the untagged sample).When an average count of 0 peptide was identified in the untagged sample, the log2FC value was instead calculated using the following formula: log2FC(average peptide count of protein *x* identified in the RNase E^1-xxx^-Strep/1).

### Bacterial adenylate cyclase two-hybrid assay

Cloned plasmids were introduced into chemically competent BTH101 cells by heat shock, and the cells were plated in LB agar supplemented with ampicillin and kanamycin. Isolated colonies were then streaked as a line of about 1-2 cm on freshly prepared Mac Conkey agar medium supplemented with ampicillin, kanamycin, 0.5 mM isopropyl β-d-thiogalactoside (IPTG), and maltose 1%. The MacConkey Petri dishes were then incubated at 30°C for 24h followed by an incubation at room temperature (∼23°C) for 24h. The bacteria were imaged by scanning the plates, after which quantification of β-galactosidase activity was done using ONPG as described in [89].

### Protein purification

His_10_Smt3-tagged PNPase, His_10_Smt3-tagged RhlB, and His_10_Smt3-tagged ArcB proteins were purified from soluble bacterial lysates by nickel-agarose affinity chromatography as described previously in the supplementary information of Hausmann *et al.* 2021 [55]. The imidazole elution profiles were monitored by SDS-PAGE. The recombinant His_10_Smt3-tagged proteins were recovered predominantly in the 250 mM imidazole fraction. Protein concentration was determined using the Bio-Rad dye reagent, with BSA as the standard, and calculated by interpolation against the BSA standard curve. The His_10_-Smt3 tag remained uncleaved in the protein-protein interaction assays.

RNase E^1-1057^, RNase E^1-588^, RNase E^1-529^, RNase E^572-1057^, RNase E^572-1057_NDPRmut^, and RNase E^572-1057_AR1mut^ proteins, each carrying a His_10_Smt3 tag at the N-terminus and a Twin-Strep-tag®-Flag at the C-terminus, were purified as described above. The 250 mM imidazole elution fractions were used directly in the Protein-Protein interaction assays. For the EMSA experiment, the RNase E proteins were further purified as follows: the 250 mM imidazole fraction were applied to a 1 ml of Strep-Tactin Sepharose (Iba) column that had been equilibrated with buffer C (50 mM Tris pH8, 200 mM NaCl, 10 % glycerol). The column was washed with 5 ml of buffer C and then eluted stepwise with 1 ml aliquots of buffer C containing, 2.5 mM D-Desthiobiotin respectively. The elution profiles were monitored by SDS-PAGE.

### Protein-protein interaction assays

Anti-Flag M2 magnetic beads (Sigma M8823) were used for the binding assay. Unless otherwise specified, a volume of 25 µL of beads suspension was used per reaction sample. The beads were washed once with 1 mL of buffer H (50 mM Tris-HCl, pH 8.0, 150 mM NaCl, 0.01% Triton X-100), before being resuspended in fresh buffer H and aliquoted into one Eppendorf tube per sample. Purified RNase E^1-529^-Flag, RNase E^1-588^-Flag RNase E^572-1057^-Flag, RNase E^572-1057_NDPRmut^-Flag, or RNase E^572-1057_AR1mut^-Flag fusion proteins (20-25 µg each) were diluted in buffer H to a final volume of 100 µL and incubated with the magnetic beads for 1 hour at 4 °C on a rotating device. When using purified RNase E^1-1057^-Flag fusion protein instead, a volume of 450 µL of purified protein (in a buffer with 50 mM Tris pH=8, 500 mM NaCl, 10% glycerol and 250 mM imidazole) was incubated with 50 µl of Anti-Flag M2 magnetic beads (Sigma M8823). After the incubation, the beads were collected using a magnetic separator and washed three times with 0.25 ml of binding buffer H to remove any unbound protein.

For affinity chromatography, the anti-Flag magnetic beads (Sigma-Aldrich) with bound RNase E-Flag fusion proteins, or unbound beads, were incubated with either 5 µg of PNPase (Figure 4A), 5 µg of RhlB (Figure 4B), or 5 µg of ArcB diluted in buffer H (final volume 100 µL, 50 µl of beads suspension used for probing ArcB binding) for 1 hour at 4 °C on a rotating device. After incubation, the beads were collected using the magnet, and the supernatant (free fraction F) was withdrawn. The anti-Flag beads were then resuspended in 100 µl of binding buffer H and subjected to three rounds of washing of 100 µl each. Following the final wash, the proteins bound to the beads were eluted with 100 µl of 2X LDS sample buffer (Invitrogen) supplemented with 167 mM DTT. Aliquots (15 µl) of the input protein sample (PNPase or RhlB), the first supernatant fraction (free fraction F), and the bead-bound fraction were mixed with 5 µl of 4X LDS sample buffer (Invitrogen), heated at 80 °C for 5 minutes, and analysed by SDS-PAGE. Polypeptides were visualized using Coomassie Blue staining.

### Growth assays on solid or liquid media

To assess the growth of the mutant strains in liquid medium, 200 μL of NYB broth was inoculated to OD_600nm_=0.05 in a 96-well plate, using an overnight culture washed twice with fresh medium. The wells on the edge of the plate were not used. The plate was incubated at 37°C using a Biotek Synergy H1 Hybrid Plate reader. OD_600nm_ values were measured every 30 minutes. Before each OD_600nm_ measurement, orbital shaking with 282 cpm (3 mm) was performed for 1 minute. Data was retrieved the following day and plotted after substraction of blank OD_600nm_ values using GraphPad Prism.

When growing strains carrying the pME6032 vector for complementation assays, NYB medium was supplemented with 50 μg/mL of Tetracycline and 70 μM of IPTG for induction. Cells were resuspended in supplemented NYB after washing and before inoculating the 96-well plate.

To assess the growth of the mutant strains on solid medium, square Petri dishes (12x12cm) were filled with 50 mL of warm NA. When testing growth at low vs high pH, either 75 μL of 37% fuming HCl or 400 μL of 5N NaOH was added per 50 mL of NA agar to achieve the desired conditions. A fresh culture (OD_600nm_= 0.4-0.8) of each strain was adjusted to OD_600nm_=1 by centrifugation and serially diluted in a 96-well plate using a multipipette. 5 μL of each dilution series was then spotted using a multipipette onto pre-dried NA plates (1 hour under a sterile flowhood). After the spots had dried, plates were incubated upside down at 37°C or 16°C.

### Galleria killing assays

*Galleria mellonella* larvae were used to assess in vivo virulence and/or survival of several RNase E mutant strains. Fresh cultures were adjusted to a suspension of OD_600nm_= 1 and serially diluted to 10^-6^in Gibco Dulbecco’s Phosphate Buffered Saline (dPBS) (Thermo Fisher Scientific). 10 μL of the 10^-6^ dilution were spotted onto NYB agar to verify that the dilutions were similar for the different strains. 10 μL spots were then put onto a clean parafilm and injected using Omnican 50 sterile insulin syringe (Braun) into the second to last leg of the larvae, facing towards the head. As a control, sterile dPBS was injected similarly into larvae to assess background levels of death. For every 10 larvae injected, the time of injection was recorded, and they were placed inside a dark incubator at 37°C. The next day, deaths were recorded every hour. A larva was considered dead when it completely stopped responding to stimuli (touched on the legs or the head with a pipette tip). Time of death was calculated as the interval of time between recorded time of injection and death occurence. Survival curves were plotted and analysed with GraphPad Prism (GraphPad Software, Prism10.2).

### RNA-seq sample collection

Independent triplicates cultures of PAO1 WT, PAO1 RNase E_1-588_, (*rne588*) and PAO1 RNase E_1-529_ (*rne529*) strains were grown in 50 mL Erlenmeyer flasks (10 mL NYB broth) until OD_600nm_= 1.0-1.2. 500 μL of each culture were immediately added into four separate 1.5 mL tubes, each containing 1 mL of RNAlater (Sigma-Aldrich, cat). After harvesting of all three strains, the tubes were centrifuged for 30 minutes at 9600 x *g*. Supernatant was discarded and pellets were stored at -80°C until RNA extraction. The RNA extraction was performed using two tubes of pellets for each strain and replicate, with the Monarch Total RNA Miniprep kit. For lysis of the bacterial cells, the two pellets were resuspended in 250 μL of 1X Tris-EDTA (TE) buffer supplemented with 1 mg/mL of lysozyme and regrouped into one tube for each strain, incubated for 5 minutes at RT, and RNA was subsequently extracted following the manufacturer’s instructions. After RNA purification, an additional DNA removal treatment was done using TURBO DNase (Thermo Fisher) according to the manufacturer’s instructions. RNA was purified again using the RNA purification columns (dark blue) from the Monarch Total RNA Miniprep kit, and following the priming, washing and elution steps described by the manufacturer. The absence of residual gDNA contamination was verified by performing a PCR for 35 cycles using primer pairs targeting *rpoD* [55]. Biological triplicate samples for each strain were prepared. Quality check, ribosomal RNA depletion and RNA sequencing were conducted by Novogene.

### RNA sequencing analysis

Reads were mapped using Bowtie2 [90] to *Pseudomonas aeruginosa* PAO1 genome assembly ASM676v1 and counts were aggregated using summarize Overlaps in the package GenomicAlignments [91] in R using the ASM676v1 gtf version 2.2 (NCBI accession: GCF_000006765.1). Differential expression was performed using DESeq2 [92] and transcripts were considered differentially expressed between two conditions if the absolute value of log2FC between conditions was greater than 1 (i.e. 2-fold up or down) with a adjusted p-value less than 0.05. To find functional enrichment among differentially expressed genes groups, the Kyoto Encyclopedia of Genes and Genomes (KEGG) was used for gene set definitions, downloaded using the package KEGGREST [93] in R. Enrichment was calculated using a hypergeometric test. All correlations presented are Pearson’s product moment correlation coefficient. Pairwise correlations of log2 Reads Per Kilobase per Million mapped reads (RPKMs) and hierarchical unsupervised clustering analysis of fold changes relative to WT demonstrate the similarity in the profile of the biological replicates for each strain. Analysis of 5’ read ends per loci was performed as described by de Araújo et al. [94] except that each dataset read number was down-sampled to 8 million.

## Acknowledgments

We thank Sylvain Guex-Crosier for his assistance with RNA extraction and Nathanaël Guggenheim for protein purification and assistance with cloning experiments. We also acknowledge the Proteomics Core Facility and the Bioinformatics Support Platform from the University of Geneva for assistance with the realisation of LC-MS-MS and bioinformatics analyses, respectively. Work in M.V. laboratory is funded by Swiss National Science Foundation (PCEFP3_203343 to M.V.) and Fondation Pierre Mercier pour la Science (to M.V.). D.G. work is supported by the Swiss National Science Foundation (PZ00P3_180142 to D.G., 2020–2023) and the Velux Foundation (grant 1814 to P.J. and D.G., 2023).

## Notes

### Competing Interest Statement

The authors have declared no competing interest.

### Summary of Updates

Supplemental files updated. Main text minor corrections.

