## Supplementary file for "Critical functions and key interactions mediated by the RNase E scaffolding domain in *Pseudomonas aeruginosa*"

**Supplementary Tables**

**Table S1: List of strains used in this study**

| Strains | Genotypes / description | Source |
|---|---|---|
| ***E. coli*** | | |
| **Rosetta (DE3)** | F- ompT hsdSB(rB- mB-) gal dcm (DE3) pRARE ($Cm^R$) | Novagen |
| **DH5α** | recA1 endA1 hsdR17 supE44 thi-1 gyrA96 relA1 Δ(lacZYA-argF)U169 [Φ80dlacZM15]F- NaIr | |
| **HB101** | proA2 hsdS20(rB- mB-) recA13 ara-14 lacYI galK2 rpsL20 supE44 xyl-5 mtl-1 F- | |
| **BTH101** | F-, cya-99, araD139, galE15, galK16, rpsL1 (Str r), hsdR2, mcrA1, mcrB1. | Euromedex |
| ***P. aeruginosa*** | | |
| **PAO1** | Wild-type | |
| **PAO1 *rne588*** | Chromosonal deletion of nucleotides encoding residues 589-1057 in *rne* | This study |
| **PAO1 *rne529*** | Chromosonal deletion of nucleotides encoding residues 530-1057 in *rne* | This study |
| **PAO1 *rne*AR1$^{mut}$** | Chromosomal arginine-to-alanine mutations within the *rne* AR1 SLiM (see Table S4) | This study |
| **PAO1 *rne*NDPR$^{mut}$** | Chromosomal point mutation of conserved residues to GGSG within the *rne* NDPR SLiM (see Table S4) | This study |
| **PAO1 *rne*AR1+AR4+REER$^{mut}$** | Chromosomal arginine-to-alanine mutations within the *rne* AR1, AR4, and REER-repeats SLiMs (see Table S4) | This study |
| **PAO1 *rne::2xStrep*** | Chromosomal insertion of a Twin Strep-tag immediately upstream of the stop codon in *rne* | This study |
| **PAO1 *rne529::2xStrep*** | Chromosonal deletion of nucleotides encoding residues 530-1057 and insertion of a Twin-Strep | This study |

| | | |
|---|---|---|
| | tag immediately upstream of the stop codon in *rne* | |
| **PAO1 *rne588::2xStrep*** | Chromosonal deletion of nucleotides encoding residues 589-1057 and insertion of a Twin-Strep tag immediately upstream of the stop codon in *rne* | This study |
| **PAO1 *rne*AR1$^{mut}$*::2xStrep*** | Chromosomal arginine-to-alanine mutations within the *rne* AR1 SLiM (see Table S4) in the strain *rne*::*2xStrep* | This study |
| **PAO1 r*ne*AR1+AR4+REER$^{mut}$*::2xStrep*** | Chromosomal arginine-to-alanine mutations within the *rne* AR1, AR4, and REER-repeats SLiMs (see Table S4) in the strain *rne*::*2xStrep* | This study |
| **PAO1 *rne::msfGFP*** | Chromosomal insertion of *msfGFP* with a flexible linker immediately upstream of the stop codon in *rne* | This study |
| **PAO1 *rne588::msfGFP*** | Chromosonal deletion of nucleotides encoding residues 589-1057 and insertion of *msfgfp* with a flexible linker immediately upstream of the stop codon in *rne* | This study |
| **PAO1 *rne529::msfGFP*** | Chromosonal deletion of nucleotides encoding residues 530-1057 and insertion of *msfgfp* with a flexible linker immediately upstream of the stop codon in *rne* | This study |
| **PAO1 *rne608::msfGFP*** | Chromosomal deletion of nucleotides encoding residues 609-1057 and insertion of msfGFP with a flexible linker immediately upstream of the stop codon in *rne* | This study |
| **PAO1 *rne733::msfGFP*** | Chromosomal deletion of nucleotides encoding residues 734-1057 and insertion of msfGFP with a flexible linker immediately upstream of the stop codon in *rne* | This study |
| **PAO1 *rne793::msfGFP*** | Chromosomal deletion of nucleotides encoding residues 794-1057 and insertion of msfGFP with a flexible linker immediately upstream of the stop codon in *rne* | This study |

| | | |
|---|---|---|
| **PAO1 *rne940::msfGFP*** | Chromosomal deletion of nucleotides encoding residues 941-1057 and insertion of msfGFP with a flexible linker immediately upstream of the stop codon in *rne* | This study |
| **PAO1 *rne*AR1$^{mut}$::*msfGFP*** | Chromosomal arginine-to-alanine mutations within the *rne* AR1 SLiM (see Table S4) in the strain *rne::msfGFP* | This study |
| **PAO1 *rne*AR4$^{mut}$::*msfGFP*** | Chromosomal arginine-to-alanine mutations within the *rne* AR4 SLiM (see Table S4) in the strain *rne::msfGFP* | This study |
| **PAO1 *rne*AR1+AR4$^{mut}$ ::*msfGFP*** | Chromosomal arginine-to-alanine mutations within the *rne* AR1 and AR4 SLiMs (see Table S4) in the strain *rne::msfGFP* | This study |
| **PAO1 *rne*REER$^{mut}$::*msfGFP*** | Chromosomal arginine-to-alanine mutations within the *rne* REER-repeats SLiM (see Table S4) in the strain *rne::msfGFP* | This study |
| **PAO1 *rne*AR1+AR4+REER$^{mut}$ ::*msfGFP*** | Chromosomal arginine-to-alanine mutations within the *rne* AR1, AR4, and REER-repeats SLiMs (see Table S4) in the strain *rne::msfGFP* | This study |
| **PAO1 *rne*NDPR$^{mut}$::*msfGFP*** | Chromosomal point mutation of conserved residues to GGSG within the *rne* NDPR SLiM (see Table S4) in the strain *rne::msfGFP* | This study |
| **PAO1 *pnp::msfGFP*/ *rne::mCherry*** | Chromosomal insertion of *msfGFP* and *mCherry* with a flexible linker immediately upstream of the stop codon in *pnp* and *rne*, respectively | This study |
| **PAO1 *rhl::msfGFP*/ *rne::mCherry*** | Chromosomal insertion of *msfGFP* and *mCherry* with a flexible linker immediately upstream of the stop codon in *rhl* and *rne*, respectively | This study |
| **PAO1 *pnp::msfGFP*** | Chromosomal insertion of *msfGFP* with a flexible linker immediately upstream of the stop codon in *pnp* | This study |
| **PAO1 *rhl::msfGFP*** | Chromosomal insertion of *msfGFP* with a flexible linker immediately upstream of the stop codon in *rhl* | This study |

| | | |
|---|---|---|
| **PAO1 *pnp::msfGFP*/ *rne529*** | Chromosomal insertion of *msfGFP* with a flexible linker immediately upstream of the stop codon in *pnp* in the strain *rne529* | This study |
| **PAO1 *rhl::msfGFP*/ *rne529*** | Chromosomal insertion of *msfGFP* with a flexible linker immediately upstream of the stop codon in *rhl* in the strain *rne529* | This study |
| **PAO1 *pnp::msfGFP*/ *rne*AR1$^{mut}$** | Chromosomal insertion of *msfGFP* with a flexible linker immediately upstream of the stop codon in *pnp* in the strain *rne*AR1mut | This study |
| **PAO1 *rhl::msfGFP*/ *rne*AR1$^{mut}$** | Chromosomal insertion of *msfGFP* with a flexible linker immediately upstream of the stop codon in *rhl* in the strain *rne*AR1mut | This study |
| **PAO1 *pnp::msfGFP*/ *rne*NDPR$^{mut}$** | Chromosomal insertion of *msfGFP* with a flexible linker immediately upstream of the stop codon in *pnp* in the strain *rne*NDPR$^{mut}$ | This study |
| **PAO1 *rhl::msfGFP*/ *rne*NDPR$^{mut}$** | Chromosomal insertion of *msfGFP* with a flexible linker immediately upstream of the stop codon in *rhl* in the strain *rne*NDPR$^{mut}$ | This study |
| **PAO1 *arcB::msfGFP*/*rne::mCherry*** | Chromosomal insertion of msfGFP and mCherry with a flexible linker immediately upstream of the stop codon in *arcB* and *rne*, respectively | This study |

**Table S2: List of plasmids used and cloning details.**

| **Name** | **Used for…** | Insert**(s) amplified with:** | **Template used in PCR** |
|---|---|---|---|
| **pME3087_RNase E1-588** | Construction of PAO1 *rne588* | p337-p338; p339-p340 | PA01gDNA |
| **pME3087_RNase E1-529** | Construction of PAO1 rne529 | p337-p341; p340-p342 | PA01gDNA |

| | | | |
|---|---|---|---|
| **pME3087_RNase E1-588-msfGFP** | Construction of PAO1 *rne588*::*msfGFP* | p365-p368; p369-p322; p323-p324 | pM13; PA01gDNA |
| **pME3087_RNase E1-529-msfGFP** | Construction of PAO1 rne529::*msfGFP* | p365-p366; p367-p322; p323-p324 | pM13; PA01gDNA |
| **pEXG2_RNase EAR1mut** | Construction of PAO1 *rneAR1mut*, PAO1 rne AR1mut::*msfGFP*, PAO1 rneAR1mut::2xStrep | p660-p661 | pSmt3_RNase E _Cter_AR1mut-SF |
| **pEXG2_RNase ENDPRmut** | Construction of PAO1 *rneNDPRmut*, *rneNDPRmut*::*msfGFP* | p662-p663; p664-p665 | pSmt3_RNase E_Cter_NDPRmut-SF; PA01 gDNA |
| **pEXG2_RNase EAR4mut** | Construction of PAO1 *rneAR4mut*::*msfGFP* | p726-p727; p728-p729; p730-p731 | pSmt3_RNase E_Cter_AR4mut-SF; PAO1 gDNA |
| **pEXG2_RNase EREERmut** | Construction of PAO1 *rneREERmut*::*msfGFP* | p720-p721; p722-p723; p724-p725 | pSmt3_RNase E_Cter_REERmut-SF; PAO1 gDNA |
| **pEXG2_RNase EAR1+AR4mut** | Construction of PAO1 *rneAR1+AR4mut*::*msfGFP* | p726-p727; p728-p729; p730-p731 | pSmt3_RNase E _Cter_AR1+AR4mut-SF; PAO1 gDNA |
| **pEXG2_RNase EAR1+AR4+REERmut** | Construction of PAO1 *rneAR1+AR4+REERmut*, PAO1 rne AR1+AR4+REERmut::*msfGFP*, PAO1 *rneAR1+AR4+REERmut*::*2xStrep* | p666-p667; p668-p669 | pSmt3_RNase E _Cter_AR1+AR4+REERmut-SF; PAO1 gDNA |
| **pME3087_RNase E-2xStrep** | Construction of PAO1 *rne::2xStrep* | p370-p379; p380-p381 | pSmt3-RNase E1-1057-SF; PAO1 gDNA |
| **pME3087_RNase E1-529-2xStrep** | Construction of PAO1 *rne529::2xStrep* strain | p378-p379; p380-p381 | pSmt3-RNase E1-529-SF; PAO1 gDNA |
| **pME3087_RNase E1-588-2xStrep** | Construction of PAO1 *rne588::2xStrep* strain | p378-p379; p380-p381 | pSmt3-RNase E1-588-SF; PAO1 gDNA |
| **pME3087_RNase E-msfGFP** | Construction of PAO1 *rne::msfGFP* strain | p319-p320; p321-p322; p323-p324 | PAO1 gDNA; msfGFP-containing plasmid |

| | | | |
|---|---|---|---|
| **pME3087_RNase E-mCherry** | Construction of PAO1 *rne::mCherry* strain | p319-p325; p326-p327; p328-p324 | PAO1 gDNA; mCherry-containing plasmid |
| **pME3087_PNPase-msfGFP** | Construction of PAO1 *pnp::msfGFP*, *pnp::msfGFP*/ *rne::mCherry*, *pnp::msfGFP* / *rne529*, *pnp::msfGFP*/*rneAR1mut*, *pnp::msfGFP*/*rneNDPRmut* strains | p405-p406; p407-p408; p409-p410 | PAO1 gDNA; msfGFP-containing plasmid |
| **pME3087_RhlB-msfGFP** | Construction of PAO1 *rhl::msfGFP, rhl::msfGFP/rne::mCherry, rhl::msfGFP/rne529*; *rhl::msfGFP/rneAR1mut*, *rhl::msfGFP/rneNDPRmut* strains | p372-p439; p441-p442; p440-p377 | PAO1 gDNA; msfGFP-containing plasmid |
| **pME3087_RNase E1-608-msfGFP** | PAO1 *rne608::msfGFP* strain | p365-p450; p455-p322; p323-p381 | PAO1 gDNA; msfGFP-containing plasmid |
| **pME3087_RNase E1-733-msfGFP** | PAO1 *rne733::msfGFP* strain | p422-p451; p456-p322; p323-p381 | PAO1 gDNA; msfGFP-containing plasmid |
| **pME3087_RNase E1-793-msfGFP** | PAO1 *rne793::msfGFP* strain | p422-p452; p457-p322; p323-p381 | PAO1 gDNA; msfGFP-containing plasmid |
| **pME3087_RNase E1-940-msfGFP** | PAO1 *rne940::msfGFP* strain | p422-p453; p458-p322; p323-p381 | PAO1 gDNA; msfGFP-containing plasmid |
| **pKT25** | BTH assay | | |
| **pKT25_PNPase** | BTH assay | p240-p241 | PAO1 gDNA |
| **pKT25_RhlB** | BTH assay | p201-p202 | PAO1 gDNA |
| **pKT25_GdhB** | BTH assay | p469-p471; p470-p472 | PAO1 gDNA |
| **pKT25_RpoB** | BTH assay | p502-p503 | PAO1 gDNA |
| **pKT25_ArcB** | BTH assay | p463-p464 | PAO1 gDNA |
| **pUT18C** | BTH assay | | |
| **pUT18C_RNase E1-1057** | BTH assay | p257-p166 | PAO1 gDNA |

| | | | |
|---|---|---|---|
| **pUT18C_RNase E1-1025** | BTH assay | p257-p438 | PAO1 gDNA |
| **pUT18C_RNase E1-940** | BTH assay | p257-p415 | PAO1 gDNA |
| **pUT18C_RNase E1-793** | BTH assay | p257-p414 | PAO1 gDNA |
| **pUT18C_RNase E1-733** | BTH assay | p257-p413 | PAO1 gDNA |
| **pUT18C_RNase E1-608** | BTH assay | p257-p412 | PAO1 gDNA |
| **pUT18C_RNase E1-588** | BTH assay | p257-p411 | PAO1 gDNA |
| **pUT18C_RNase E1-529** | BTH assay | p257-p295 | PAO1 gDNA |
| **pUT18C_RNase E940-1057** | BTH assay | p449-p166 | PAO1 gDNA |
| **pUT18C_RNase E733-793** | BTH assay | p165-p414 | PAO1 gDNA |
| **pUT18C_RNase E733-761** | BTH assay | p165-p590 | PAO1 gDNA |
| **pUT18C_RNase E762-793** | BTH assay | p589-p414 | PAO1 gDNA |
| **pME6032** | | | |
| **pME6032_RNase E** | Complementation assay | p403-p790 | pME6032_RNase E-msfGFP |
| **pME6032_RNase E-msfGFP** | Complementation assay | p403-p404? | PAO1 *rne::msfGFP* gDNA |
| **pME6032_RNase E-mCherry** | Complementation assay | p403-p791 | pSmt3_RNase E-mCherry |
| **pSmt3_RNase E1-529-SF** | Protein purification | p181-p185 | PAO1 gDNA |
| **pSmt3_RNase E1-588-SF** | Protein purification | p181-p396 | PAO1 gDNA |
| **pSmt3_RNase E572-1057-SF** | Protein purification | p182-481 | PAO1 gDNA |
| **pSmt3_RNase E572-1057-$^{AR4mut-SF}$** | Protein purification | p555-p557; p558-p559; p560-p561; p564-p565; p562-p563; p566-p568 | PA01 gDNA; AR4mut |

| | | | |
|---|---|---|---|
| **pSmt3_RNase E572-1057–$^{\text{AR1mut-SF}}$** | Protein purification | p555-p557; p558-p559; p560-p561; p564-p565; p562-p563; p566-p568 | PA01 gDNA; AR1mut |
| **pSmt3_RNase E572-1057–$^{\text{REERmut-SF}}$** | Protein purification | p555-p557; p558-p559; p560-p561; p564-p565; p562-p563; p566-p568 | PA01 gDNA; REERmut |
| **pSmt3_RNase E572-1057–$^{\text{AR1+AR4mut-SF}}$** | Protein purification | p555-p557; p558-p559; p560-p561; p564-p565; p562-p563; p566-p568 | PA01 gDNA; AR4mut; AR1mut |
| **pSmt3_RNase E572-1057–$^{\text{AR1+AR4+REERmut-SF}}$** | Protein purification | p555-p557; p558-p559; p560-p561; p564-p565; p562-p563; p566-p568 | PA01 gDNA; AR4mut; AR1mut ; REERmut |
| **pSmt3_RNase E572-1057–$^{\text{NDPRmut-SF}}$** | Protein purification | p555-p557; p558-p559; p560-p561; p564-p565; p562-p563; p566-p568 | PA01 gDNA; NDPR |
| **pSmt3-PNPase** | Protein purification | p244-p245 | PAO1 gDNA |
| **pSmt3-RhlB** | Protein purification | p498-p499 | PAO1 gDNA |
| **pEXG2_ArcB-msfGFP** | Construction of PAO1 *arcB::msfGFP/rne::mCherry* | p828-p829, p830-p831, p832-p833 | PAO1 gDNA; msfGFP-containing plasmid |
| **pSmt3-ArcB** | Protein purification | p834-p835 | |

**Table S3: List of primers used in this study.**

| Primer code | Primer sequence (5' -> 3') |
|---|---|
| p165 | GTACGGATCCCGCCCTGGAAGCCGAGGCATTG |
| p166 | GATCGAATTCTCAGGCATGCGGGCGCG |
| p181 | GTACGGATCCAAAAGAATGCTGATCAACGCGACTC |
| p182 | CTAGGCGGCCGCGGCATGCGGGCGCGA |
| p185 | CTAGGCGGCCGCAGCGACGGTCTTGACTGC |
| p201 | GTACGGATCCCCTCAAAGCACTCAAGAAAATCTTCG |
| p202 | GATCGAATTCTCAGTGCTTGCGCGGTAC |
| p240 | GTACGGATCCCAACCCGGTAACCAAGCAATTC |
| p241 | GATCGAATTCTTACACGCCGGAAGCTTC |
| p244 | CAGTGGATCCGGAGGAAGCAACCCGGTAACCAAGCAATTC |
| p245 | CATGGAGCTCTTACACGCCGGAAGCTTCG |
| p257 | GTACGGATCCCAAAAGAATGCTGATCAACGCGACTC |
| p295 | GATCGAATTCTCAAGCGACGGTCTTGACTGC |
| p319 | TAGGAGCATAGAATTCGCAAGCCACTGAAGAAGCTCC |
| p320 | CTTCACCTTTACGGGAACCACCACCGGCATGCGGGCGCGA |
| p321 | CGCCCGCATGCCGGTGGTGGTTCCCGTAAAGGTGAAGAACTG |
| p322 | CCTTTTTTCAGCCTGTCACTTATTTGTAGAGTTCATCCA |
| p323 | GATGAACTCTACAAATAAGTGACAGGCTGAAAAAAGG |
| p324 | GAATGGCAAAAGCTTCTTCAAGAACCCGCTGGTG |
| p325 | CTCCTCGCCCTTGCTCACGGAACCACCACCGGCATGCGGGCGCGA |
| p326 | CGCCCGCATGCCGGTGGTGGTTCCGTGAGCAAGGGCGAGGAG |
| p327 | CCTTTTTTCAGCCTGTCACTTACTTGTACAGCTCGTCCATG |
| p328 | CATGGACGAGCTGTACAAGTAAGTGACAGGCTGAAAAAAGG |
| p337 | TAGGAGCATAGAATTCCGACGTCGAGTCGCTGTC |
| p338 | TCAGCCTGTCACTCAGCGGGTTTGGCGCTCG |
| p339 | GCCAAACCCGCTGAGTGACAGGCTGAAAAAAGG |
| p340 | GAATGGCAAAAGCTTGCAGCTTCTTCAAGAACC |
| p341 | CAGCCTGTCACTCAAGCGACGGTCTTGACTGC |
| p342 | CAAGACCGTCGCTTGAGTGACAGGCTGAAAAAAGG |
| p365 | TAGGAGCATAGAATTCCCTGATCGTCATCGACTTCATCG |
| p366 | CAGTTCTTCACCTTTACGGGAACCACCACCAGCGACGGTCTTGACTGC |
| p367 | CAAGACCGTCGCTGGTGGTGGTTCCCGTAAAGGTGAAGAACTG |
| p368 | CAGTTCTTCACCTTTACGGGAACCACCACCGCGGGTTTGGCGCTCG |
| p369 | GCCAAACCCGCGGTGGTGGTTCCCGTAAAGGTGAAGAACTG |
| p370 | TAGGAGCATAGAATTCGAGCTGGAAGGCAGCGAG |
| p372 | TAGGAGCATAGAATTCGAGGTGCACCTGGACATGG |

| | |
|---|---|
| **p377** | GAATGGCAAAAGCTTGCATCGCAGAGCAGCAAGG |
| **p378** | TAGGAGCATAGAATTCGCAGACCAACCTGGAAGC |
| **p379** | CCTTTTTTCAGCCTGTCACTCATCCGCTAGCTCCTTTCTCG |
| **p380** | GGAGCTAGCGGATGAGTGACAGGCTGAAAAAAGG |
| **p381** | GAATGGCAAAAGCTTGCAGCTTCTTCAAGAACC |
| **p396** | CTAGGCGGCCGCGCGGGTTTGGCGCTCG |
| **p403** | GATCGAATTCATGAAAAGAATGCTGATCAAC |
| **p404** | GATCGGTACCTTATTTGTAGAGTTCATCCATG |
| **p405** | CAATATAGGAGCATAGAATTCGACATCAAGATCAACGGTATCAC |
| **p406** | CCTTTACGGGAACCACCACCCACGCCGGAAGCTTCG |
| **p407** | CGAAGCTTCCGGCGTGGGTGGTGGTTCCCGTAAAGGTGAAGAACTG |
| **p408** | TTCATCGATCACGACTCGACTTATTTGTAGAGTTCATCCA |
| **p409** | TGGATGAACTCTACAAATAAGTCGAGTCGTGATCGATGAAAAC |
| **p410** | GAGAATGGCAAAAGCTTGGCTTTCACGCCCTTGATG |
| **p411** | GATCGAATTCTCAGCGGGTTTGGCGCTCGG |
| **p412** | GATCGAATTCTCAATTGCCATCGCGGCCATC |
| **p413** | GATCGAATTCTCAGGCCGCGGCGTCGCGAG |
| **p414** | GATCGAATTCTCACGCGGCGGCGTTATCGGTC |
| **p415** | GATCGAATTCTCAGATGGCCGGGGTCGGCTC |
| **p422** | TAGGAGCATAGAATTCGCAGTCAAGACCGTCGCTC |
| **p438** | GATCGAATTCTCATTCCTGGGCTTCGTCCGCCTG |
| **p439** | CCTTTACGGGAACCACCACCGTGCTTGCGCGGTACCGGCT |
| **p440** | TGGATGAACTCTACAAATAAACGCAGGCATGGAGAA |
| **p441** | AGCCGGTACCGCGCAAGCACGGTGGTGGTTCCCGTAAAGGTGAAGAACTG |
| **p442** | TTTTTCTCCATGCCTGCGTTTATTTGTAGAGTTCATCCA |
| **p449** | GTACGGATCCCGAGCCGACCCCGGCCATC |
| **p450** | CAGTTCTTCACCTTTACGGGAACCACCACCATTGCCATCGCGGCCATC |
| **p451** | CAGTTCTTCACCTTTACGGGAACCACCACCGGCCGCGGCGTCGCGAG |
| **p452** | CAGTTCTTCACCTTTACGGGAACCACCACCCGCGGCGGCGTTATCGGTC |
| **p453** | CAGTTCTTCACCTTTACGGGAACCACCACCGATGGCCGGGGTCGGCTC |
| **p455** | GATGGCCGCGATGGCAATGGTGGTGGTTCCCGTAAAGGTGAAGAACTG |
| **p456** | CTCGCGACGCCGCGGCCGGTGGTGGTTCCCGTAAAGGTGAAGAACTG |
| **p457** | GACCGATAACGCCGCCGCGGGTGGTGGTTCCCGTAAAGGTGAAGAACTG |
| **p458** | GAGCCGACCCCGGCCATCGGTGGTGGTTCCCGTAAAGGTGAAGAACTG |
| **p463** | GTACAGATCTCGCTTTCAACATGCACAACC |
| **p464** | GATCGAATTCTCAGATGTCGGCGAGG |
| **p469** | CTGCAGGGTCGACTCTAGAGGCGTTCTTCACCGCG |
| **p470** | GTACGAGATCTTCGTCTACTC |
| **p471** | GAGTAGACGAAGATCTCGTAC |
| **p472** | ACGTTGTAAAACGACGGCCGTCAGGGGATGCAGACAC |

| | |
|---|---|
| **p481** | GTACGGATCCGCCGCCAAGCCTGCTGAAAC |
| **p498** | CAGTAAGCTTCAGTGCTCAAAGCACTCAAGAAAATC |
| **p499** | CATGCTCGAGTCAGTGCTTGCGCGGTAC |
| **p502** | GTACGGATCCCATGGCTTACTCATACACTG |
| **p503** | GATCCAATTGTTATTCGGTTTCCAGTTCGAT |
| **p555** | GGATCCGCCGCCAAGCCTGCTGAAAC |
| **p557** | CTCTTCGTCGCGGCGAT |
| **p558** | GGCAATCGCCGCGAC |
| **p559** | TGCCTCGGCTTCCAGGG |
| **p560** | GACGCCGCGGCCC |
| **p561** | CGCGGCGGCGTTATCGG |
| **p562** | CTGCTGGTCGCGCAAG |
| **p563** | GGCCGGCACCGCC |
| **p564** | GGCAGCGAGGCGACC |
| **p565** | AGCTTCTTCAGTGGCTTGCG |
| **p566** | GAGGAGCCGACCCCGG |
| **p568** | GCGGCCGCGGCATGCGGGCGCGA |
| **p589** | GTACGGATCCCTCCCGCGGCCAGCGTCGT |
| **p590** | GATCGAATTCTCAGCGGCGGCGCGGGCGCTC |
| **p660** | GTCGACTCTAGAGGATCCGCCGCCAAGCCTGCTGA |
| **p661** | GAAATTAATTAAGGTACCGAATTCCGCTTCGGCTGCTGCT |
| **p662** | GTCGACTCTAGAGGATCCCCGAACGACGAGAGCCT |
| **p663** | TCGACGGCGGGAGCGGCCTGAGC |
| **p664** | CGCTCCCGCCGTCGAGGAGATCCCG |
| **p665** | GAAATTAATTAAGGTACCGAATTCGCTCATCGACAAGGGCGG |
| **p666** | GTCGACTCTAGAGGATCCGGCATCGTCTGCCCGC |
| **p667** | TTGGCGGCAGGTTGATCCTTGCCTGCG |
| **p668** | TCAACCTGCCGCCAAGCCTGC |
| **p669** | GAAATTAATTAAGGTACCGAATTCCTCGACGGCGGGAGCG |
| **p720** | GTCGACTCTAGAGGATCCGGAAGAAGCCCTGAAGGACCG |
| **p721** | CGATTGCCATCGCGGCCATCGCG |
| **p722** | CCGCGATGGCAATCGCCGCGAC |
| **p723** | TTCGGCAATGCCTCGGCTTCCAGGG |
| **p724** | CGAGGCATTGCCGAACGACGAGAGCC |
| **p725** | GAAATTAATTAAGGTACCGAATTCCGTTGCCGGTTGCACG |
| **p726** | GTCGACTCTAGAGGATCCGACGTCGAGTCGCTGTCG |
| **p727** | GCTTGCTGGTTTCAGCAGGCTTGGCG |
| **p728** | GCTGAAACCAGCAAGCCGGCTGC |
| **p729** | AGCGGCGCGGCGGCGTTATCGGTCG |
| **p730** | CGCCGCCGCGCCGCTGAACACC |

| | |
|---|---|
| **p731** | **GAAATTAATTAAGGTACCGAATTCCGTTGCAGGCGACGTTTTT** |
| **p790** | **GATCGGTACCTCAGGCATGCGGGCGC** |
| **p791** | **GATCGGTACCCTACTTGTACAGCTCGTCCATG** |

**Table S4: Sequence of mutated SLiMs**

| RNase E SLiM | Sequence |
|---|---|
| **AR4** | RQTRQDERRNGRQQNRRRDGRDGNRRDEE |
| **AR4mut** | RQTAADEAANGAAQNAAADGADGNRRDEE |
| **REER** | RKPREERAERQPREERAERPNREERSERRREERAERPAREERQPREGREERAERTPREERQPREGREGREERSERRREERAERPAREERQPREGREERAERPAREERQPREDRQARDAA |
| **REERmut** | AKPAEEAAEAQPAEEAAEAPNAEEASEAAAEEAAEAPAAEEAQPAEGAEEAAEATPAEEAQPAEGAEGAEEASEAAAEEAAEAPAAEEAQPAEGAEERAEAPAAEEAQPAEDAQAADAA |
| **AR1** | GERPRRRSRGQRRRSNRRERQREVSGELEGSEATDNAA |
| **AR1mut** | GEAPAAASAGQAAASNAAEAQAEVSGELEGSEATDNAA |
| **NDPR** | GRALNDPREKRRLQREAERLAREAAAAAEAAAQAA |
| **NDPRmut** | GRALGGSGEKRRLQRGGSGLAREGGSGAEAAAQAA |

**Table S5**: **RNAseq data (see Excel file).**

Sheet 1. Comparisons rne529 vs rne588 targets (_over WT)

Sheet 2. Differentially regulated sRNAs genes

Sheet 3: KEGG enrichment: gene downregulated in the rne529 strain vs WT

Sheet 4: KEGG enrichment: gene upregulated in the rne529 strain vs WT

Sheet 5: KEGG enrichment: gene downregulated in the rne588 strain vs WT

Sheet 6: KEGG enrichment: gene upregulated in the rne588 strain vs WT

#### Supplementary Figures Legends

**Figure S1**: **Alignment of *E. coli* (Ec) and *P. aeruginosa* (Pa) RNase E orthologs**. Alignment of the Ec rne gene with the Pa homolog (PA2976) was done using the EBLOSUM62 matrix.

**Figure S2**: **Variability of RNase E SLiMs between *P. aeruginosa* and *E. coli* or other Pseudomonas species.** **(A)** Comparison of *E. coli* (Ec) and *P. aeruginosa* (Pa) RNase E SLiMs based on the analysis performed in [1]. The precise amino acid positions were determined by manual sequence inspection. **(B)** Analysis of the frequency of RE[ED]R repeats within the *Pseudomonas* genus (see methods for details).

**Figure S3**: **Logo plot displaying the consensus sequence of global alignment of RNase E orthologs within the Pseudomonas group.** 5000 RNase E ortholog sequences were downloaded from the NCBI database, aligned using Clustal Omega and created using Weblogo [2]. Identified SLiMs are shown within boxes of different colours and all overlap with SLiMs previously identified in [1], though they are typically longer than previously described SLiMs. Position of the SLiMs on the RNase E protein of the *P. aeruginosa* PAO1 strain is indicated (see also Fig 1B).

**Figure S4**: **RNase E CTD RNA binding: extended EMSA. (A-B)** EMSA with purified RNase E C-terminal domain (residues 572-1057) having either the native sequence (WT) or systematic alanine mutation of arginine residues within the REER-rich region (REERmut) (B), the AR4 SLiM (AR4mut) (A) or the AR1 SLiM (AR1mut) (A). The *malEF* mRNA was used as a substrate. Ratios of RNA:protein ranging from 1:0.3 to 1:10 (A) or 1:0.3 to 1:80 (B) were tested. **(C)** Quantification of average signal intensity in PAO1 strains shown in Figure 2B. I: RNase E::msfgfp, II: RNase $E^{1-588}$::msfGFP III: RNase $E^{1-529}$::msfGFP, IV: RNase $E^{AR4mut}$::msfGFP, V: RNase $E^{REERmut}$::msfGFP, VI: RNase $E^{AR1mut}$::msfGFP, VII: RNase $E^{AR1+AR4mut}$::msfGFP, VIII: RNase $E^{AR1+AR4+REERmut}$::msfGFP. Fluorescence intensity was measured along the longitudinal axis of the cells in 50-100 cells using Fiji. The intensity values were corrected by subtraction of background noise values. Quantification of the variance of pixel intensity normalised by the average cell intensity was done using R. Statistical significance was assessed by performing t-tests using R. p-values for comparisons tested are indicated as follows: *:<0.05; **:<0.01; ***:<0.001; ****:<0.0001, n.s.: not significant (95% confidence interval). **(D)** Western Blot showing expression levels of chromosomal Strep-tagged RNase E variants. 1: WT RNase E sequence, 2: RNase $E^{1-588}$ truncation, 3: RNase $E^{1-529}$ truncation, 4: RNase $E^{AR1mut}$ point mutant 5: RNase $E^{AR1+AR4+REERmut}$ point mutant. As a loading control, the expression levels of the major outer membrane protein OprF were probed using anti-OprF antibodies.
**(E)** Representative images of live PAO1 strains expressing truncated variants of the RNase E::msfGFP fusion as described above each image. The scale is 2 μm.

**(F)** Pooled histograms showing the distribution of the measured signal intensity corrected by the mean signal intensity in the corresponding cell. Representative images of each RNase E-msfGFP variant or exposure conditions are shown in Figure 2 B,D.

**Figure S5**: **Pull-down eluates analysis and direct binding assay using purified ArcB protein. (A)** Silver staining of pull-down eluates obtained using RNase $E^{1\text{-}1057}$-Strep or RNase $E^{1\text{-}529}$-Strep as a bait protein. The negative control was done using cellular lysates from the WT strain (untagged). Direct resuspension of the MagStrep® Strep-Tactin®XT magnetic beads in 1x Laemmli buffer was done. kDa: kiloDaltons. The ProteoSilver Silver Stain kit (Sigma) was used according to the manufacturer's instructions. **(B)** Direct protein-protein binding assay between PNPase and RNase $E^{1\text{-}588}$ . I: input PNPase fraction, F: free PNPase fraction, B: bound PNPase fraction. **(C)** Direct protein-protein binding assay between ArcB and RNase $E^{1\text{-}1057}$ or RNase $E^{1\text{-}588}$. I: input fractions as indicated, F: free ArcB fraction, B: bound ArcB fraction. The binding assay was performed twice and a representative gel is shown. For **(B-C)**, purified His-Smt3-RNase $E^{1\text{-}1057}$-Strep-FLAG or His-Smt3-RNase $E^{1\text{-}588}$-Strep-FLAG proteins were used as baits, immobilized to anti-flag magnetic beads and incubated with purified His-Smt3-PNPase or His-Smt3-ArcB proteins as indicated to assess direct protein binding.

**Figure S6: Bacterial two hybrid (BTH) results.** Visualisation of the bacterial two-hybrid *E. coli* BTH101 patches used for quantification of β-galactosidase activity (Figure 3 C-F). Isolated colonies were patched onto McConkey agar, allowing visualisation of a red pigmentation if the bacteria are able to metabolise maltose and lactose due to restored cAMP production. **(A)** *E. coli* BTH101 co-expressing T18-RNase E with one of five T25-fused putative protein partners, as indicated below each picture. **(B-D)** *E. coli* BTH101 co-expressing T25-PNPase **(B)**, T25-RhlB **(C)**, or T25-ArcB **(D)** with either T18-RNase E or one of eight to ten truncated T18-RNase E fragments, as indicated below each picture.

**Figure S7**: **Fitness of chromosomally-tagged strains and subcellular localisation of RNase E and/or ArcB. (A)** Growth curves of PAO1 strains expressing chromosomal msfGFP-tagged or mCherry-tagged versions of RNase E (*rne::msfGFP* / *rne::mCherry*), PNPase (*pnp::msfGFP*), or RhlB (*rhl::msfGFP*). Bacteria were cultivated in NYB medium at 37°C in a 96-well plate (200 µL per well), with a shaking of 1 minute at 282 cpm (3 mm amplitude) before each $OD_{600nm}$ reading. All the strains tested grow nearly identically. **(B-C)** Representative images of a strain expressing RNase $E^{NDPRmut}$::msfGFP **(B)**, highlighting the formation of foci similar to those observed for RNase E::msfGFP (shown in Figure 2), or a strain co-expressing ArcB-msfGFP and RNase E-mCherry **(C)**, showing a completely smooth subcellular localisation for ArcB-msfGFP rather than a foci localisation which is expected for a core RNA degradosome component. Scale is 2 µm. The images were corrected for background noise as described in Methods.

**Figure S8. Complementation assay.** Expression of RNase E, RNase E-msfGFP or RNase E-mCherry from the IPTG-inducible pME6032 vector can complement the growth delay of *rne529* or *rne*AR1+AR4+REER$^{mut}$ chromosomal mutant strains at 37°C. For reference, the WT strain was also transformed with the pME6032 vectors. For growth details, see Materials and Methods.

**Figure S9: RNA sequencing: quality control and assessment of similarity of replicates and comparison of strains. (A)** Pairwise Pearson correlation of log10 RPKMs across all samples. **(B)** Unsupervised hierarchical clustering of log10 RPKMs using a Euclidiean distance measure.

**Figure S10: Histogram of ratio of coverage in each sample compared to each WT sample across all genomic positions.** Each replicate of every strain is compared, pairwise with each WT replicate. For each genomic position, a ratio is calculated: the coverage of 5'ends of reads in the first sample is divided by the total coverage across this sample and WT replicate (all samples are downsampled to 8 million reads). A histogram of the frequencies of these ratios is plotted. The primary focus is whether we see an enrichment of ratios of 1 in the deletion, indicating an increase in positions solely with 5'end coverage in the deletion, suggesting potential cleavage points not present in WT.

### Figure S1

```
Aligned_sequences: 2
# 1: RNE_ECOLI
# 2: PA2976
# Matrix: EBLOSUM62
# Gap_penalty: 10.0
# Extend_penalty: 0.5
#
# Length: 1167
# Identity:     530/1167 (45.4%)
# Similarity:   651/1167 (55.8%)
# Gaps:         216/1167 (18.5%)
# Score: 2133.0
#
Domains: NTER, MTS, AR, RhlB binding site (Ec), Enolase binding (Ec), AEPV rich, PNPase
binding site (Ec), REERs (Pa),NDPR (Pa)
# truncations
#=======================================

RNE_ECOLI          1 MKRMLINATQQEELRVALVDGQRLYDLDIESPGHEQKKANIYKGKITRIE     50
                     ||||||||||.|||||||||||||:||||||...||||||||||:|||:|
PA2976             1 MKRMLINATQPEELRVALVDGQRLFDLDIESGAREQKKANIYKGRITRVE     50

RNE_ECOLI         51 PSLEAAFVDYGAERHGFLPLKEIAREYFPANYSAHGRPNIKDVLREGQEV    100
                     |||||||||:|||||||||||||:||||  ..|..||.|||:||.|||||
PA2976            51 PSLEAAFVDFGAERHGFLPLKEISREYF--KKSPEGRINIKEVLSEGQEV     98

RNE_ECOLI        101 IVQIDKEERGNKGAALTTFISLAGSYLVLMPNNPRAGGISRRIEGDDRTE    150
                     |||::|||||||||||||||||||.||||||||||||||||||||::|.|
PA2976            99 IVQVEKEERGNKGAALTTFISLAGRYLVLMPNNPRAGGISRRIEGEERNE    148

RNE_ECOLI        151 LKEALASLELPEGMGLIVRTAGVGKSAEALQWDLSFRLKHWEAIKKAAES    200
                     |:|||..|..|..|||||||||:|:|.|.|||||.:.|:.|.|||:|:..
PA2976           149 LREALNGLNAPADMGLIVRTAGLGRSTEELQWDLDYLLQLWSAIKEASGE    198

RNE_ECOLI        201 RPAPFLIHQESNVIVRAFRDYLRQDIGEILIDNPKVLELARQHIAALGRP    250
                     |.|||||:||||||:||.||||||||||:|||:....|.|...|..: .|
PA2976           199 RGAPFLIYQESNVIIRAIRDYLRQDIGEVLIDSIDAQEEALNFIRQV-MP    247

RNE_ECOLI        251 DFSSKIKLYTGEIPLFSHYQIESQIESAFQREVRLPSGGSIVIDSTEALT    300
                     .::||:|||...:|||:.:|||||||:||||||:||||||||||.||||.
PA2976           248 QYASKVKLYQDSVPLFNRFQIESQIETAFQREVKLPSGGSIVIDPTEALV    297

RNE_ECOLI        301 AIDINSARATRGGDIEETAFNTNLEAADEIARQLRLRDLGGLIVIDFIDM    350
                     :|||||||||:||||||||..||||||:||||||||||:|||||||||||
PA2976           298 SIDINSARATKGGDIEETALQTNLEAAEEIARQLRLRDIGGLIVIDFIDM    347

RNE_ECOLI        351 TPVRHQRAVENRLREAVRQDRARIQISHISRFGLLEMSRQRLSPSLGESS    400
                     ||.::|||||.|:|||:..||||:|:..||||||||||||||.|||||:|
PA2976           348 TPAKNQRAVEERVREALEADRARVQVGRISRFGLLEMSRQRLRPSLGETS    397

RNE_ECOLI        401 HHVCPRCSGTGTVRDNESLSLSILRLIEEEALKENTQEVHAIVPVPIASY    450
                     ..|||||:|.|.:||.|||||:|||||||||||:.|.||.|.||..:|::
PA2976           398 GIVCPRCNGQGIIRDVESLSLAILRLIEEEALKDRTAEVRARVPFQVAAF    447

RNE_ECOLI        451 LLNEKRSAVNAIETRQDGVRCVIVPNDQMETPHYHVLRVRKGEETPTL--    498
                     ||||||:|:..||.| ...|..|:|:|.:||||:.|.|:|  :::|.|
PA2976           448 LLNEKRNAITKIELR-TRARIFILPDDHLETPHFEVQRLR--DDSPELVA    494

RNE_ECOLI        499 ---SYMLPKL-HEEAMALPSEEEFAERKRPEQPALATFAMPDVPPAPTPA    544
                        ||.:..: ||||..:.|......    ::.|:.|.|         |.
PA2976           495 GQTSYEMATVEHEEAQPVSSTRTLVR----QEAAVKTVA---------PQ    531

RNE_ECOLI        545 EPAAPVVAPAPKAAPATPAAP-AQPGLLSRFFGALKALFSGGEE--TKPT    591
                     :|     ||....||..||.| .:|.|......:|..||:|.::  .||.
PA2976           532 QP-----APQHTEAPVEPAKPMPEPSLFQGLVKSLVGLFAGKDQPAAKPA    576
```

```
RNE_ECOLI        592 EQPAPKAEAKPERQQDRRKPRQNNRR-------DRNERRDTRSERTEGS-     633
                     |...|.|| :..||.:||..||.|||       .|:|.|..|.||.|..
PA2976           577 ETSKPAAE-RQTRQDERRNGRQQNRRRDGRDGNRRDEERKPREERAERQP     625

RNE_ECOLI        634 -------DNREENRRNRRQAQQQTAETRESRQQAEVTEK--ARTADEQQA     674
                            .|||| |..||:.::.....||.||..|..|:  .||..|::.
PA2976           626 REERAERPNREE-RSERRREERAERPAREERQPREGREERAERTPREERQ     674

RNE_ECOLI        675 PRRERSRRRNDDKRQAQQEAKALNVEEQSVQETEQEERVRPVQPRRKQRQ     724
                     ||..|..|....:|:.::.|:....||:..:|..:|...||.:..|:.|
PA2976           675 PREGREGREERSERRREERAERPAREERQPREGREERAERPAREERQPR-     723

RNE_ECOLI        725 LNQKVRYEQSVAEEAVVAPVVEETVAAEPIVQEAPA-PRTELVKVPLPVV     773
                                          |:..|.:....||.| |..|.:
PA2976           724 ---------------------EDRQARDAAALEAEALPNDESL-------     745

RNE_ECOLI        774 AQTAPEQQEENNADNRDNGGMPRRSRRSPRHLRVSGQRRRRYRDERY---     820
                          ||.|:::.|    |..|||..|        ||||||..|.||.
PA2976           746 -----EQDEQDDTD----GERPRRRSR--------GQRRRSNRRERQREV     778

RNE_ECOLI        821 ---------PTQSPMPL-TVACASPELASGKVWIRYPIVRPQDVQVEEQR     860
                              ...:..|| |||.|:   |:|             :.|..:.
PA2976           779 SGELEGSEATDNAAAPLNTVAAAA---AAG-------------IAVASEA     812

RNE_ECOLI        861 EQEEVHVQPMVTEVPVA---AAIEPVVSAPVVEEVAGV-----------V     896
                     .:..|...|..|....:   |:.|...|.....|..|.           :
PA2976           813 VEANVEQAPATTSEAASETTASDETDASTSEAVETQGADSEANTGETADI     862

RNE_ECOLI        897 EAPVQVAEPQPEVVETTHPEVIAAAVTEQPQVITESDVAVAQEVAEQAEP     946
                     ||||.|:..:.|..::|   ::.|..||:....:||     .|..|.||.
PA2976           863 EAPVTVSVVRDEADQST---LLVAQATEEAPFASES-----VESREDAES     904

RNE_ECOLI        947 VVEPQEETADIEEVVETAEVVVAEPE-----------VVAQPA-------     978
                     .|:|..|.|  |||.....|.||.|.           :.|.||
PA2976           905 AVQPATEAA--EEVAAPVPVEVAAPSEPAATEEPTPAIAAVPANATGRAL     952

RNE_ECOLI        979 -------------------APVVAEVAAE----VETVAAVEPEVTVEHNH    1005
                                        |...||.||:    ||.:.||..|.......
PA2976           953 NDPREKRRLQREAERLAREAAAAAEAAAQAAPAVEEIPAVASEEASAQEE    1002

RNE_ECOLI       1006 ATAP----MTRAPAPEYVPEA-------PRHSDWQRPTFAFEGKGAAGGH    1044
                     ..||    :|:|..|....||       |..|          |:|||...
PA2976          1003 PAAPQAEEITQADVPSQADEAQEAVQAEPEAS----------GEGAADTE    1042

RNE_ECOLI       1045 TATHHASAAPARPQPVE   1061
                     .|.....:..:||..
PA2976          1043 HAKKTEESETSRPHA--   1057
```

### Figure S2

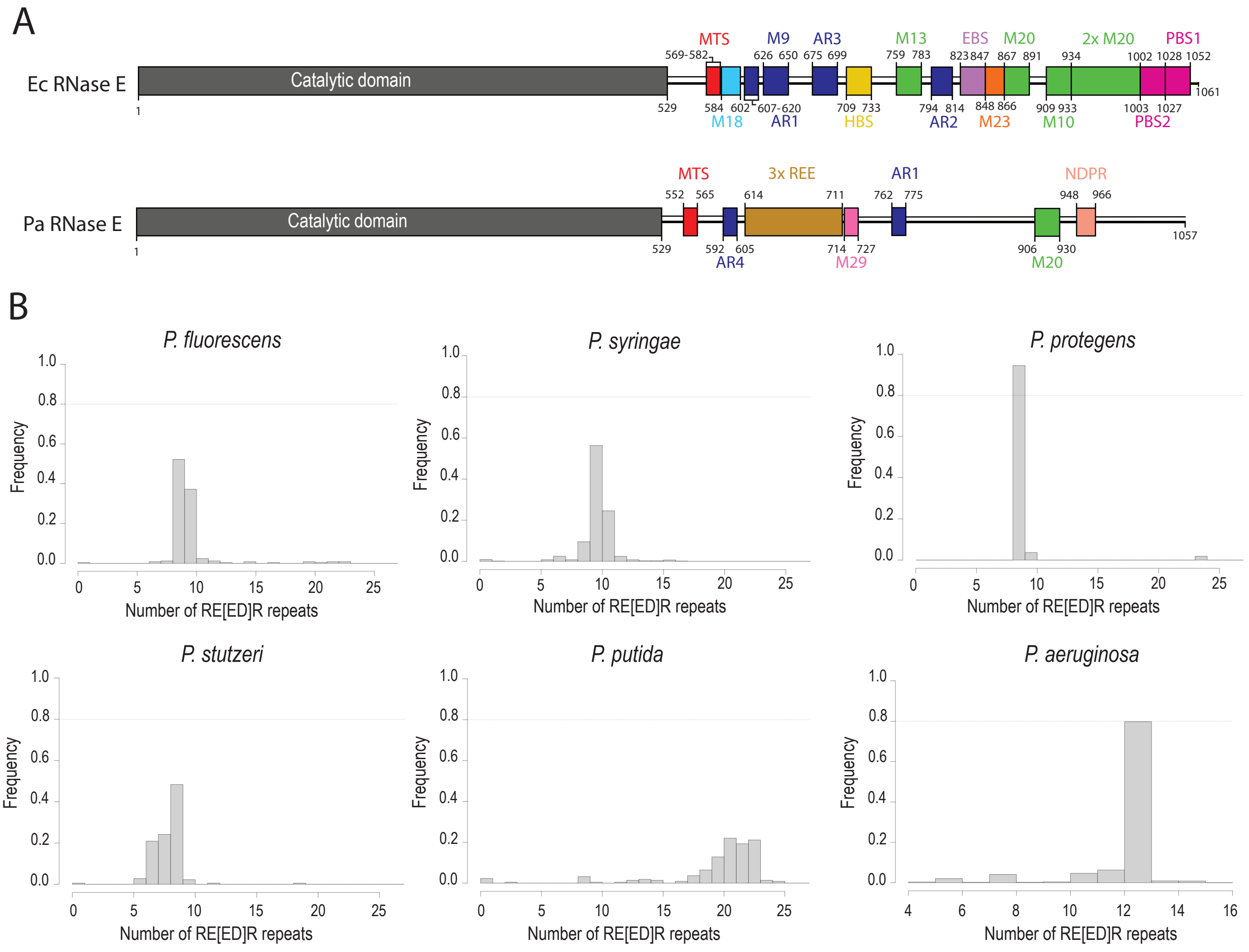

A
Ec RNase E
Catalytic domain
MTS
M9
AR3
M13
EBS
M20
2x M20
PBS1
M18
AR1
HBS
AR2
M23
M10
PBS2
Pa RNase E
Catalytic domain
MTS
3x REE
AR1
NDPR
AR4
M29
M20
B
P. fluorescens
P. syringae
P. protegens
P. stutzeri
P. putida
P. aeruginosa
Frequency
Number of RE[ED]R repeats

Figure S3

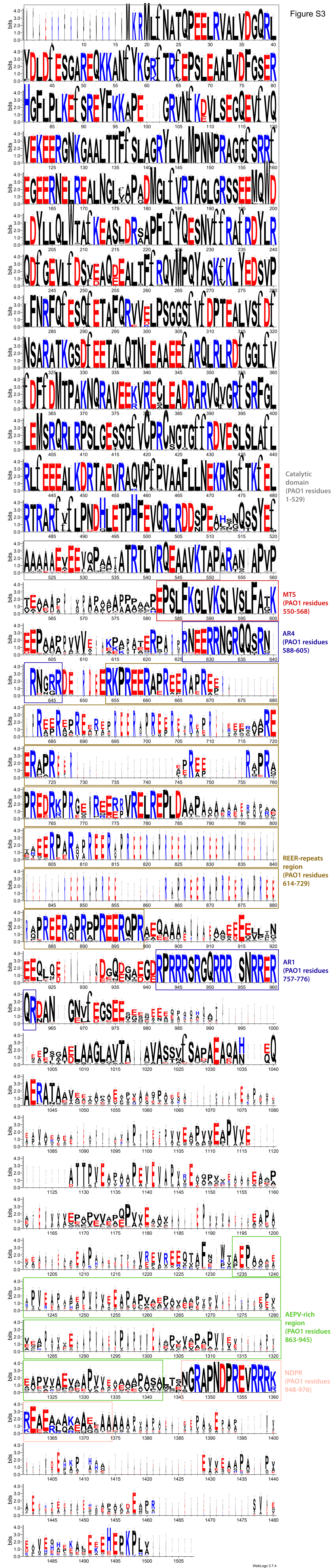

Catalytic domain (PAO1 residues 1-529)
MTS (PAO1 residues 550-568)
AR4 (PAO1 residues 588-605)
REER-repeats region (PAO1 residues 614-729)
AR1 (PAO1 residues 757-776)
AEPV-rich region (PAO1 residues 863-945)
NDPR (PAO1 residues 948-976)
WebLogo 3.7.4

### Figure S4

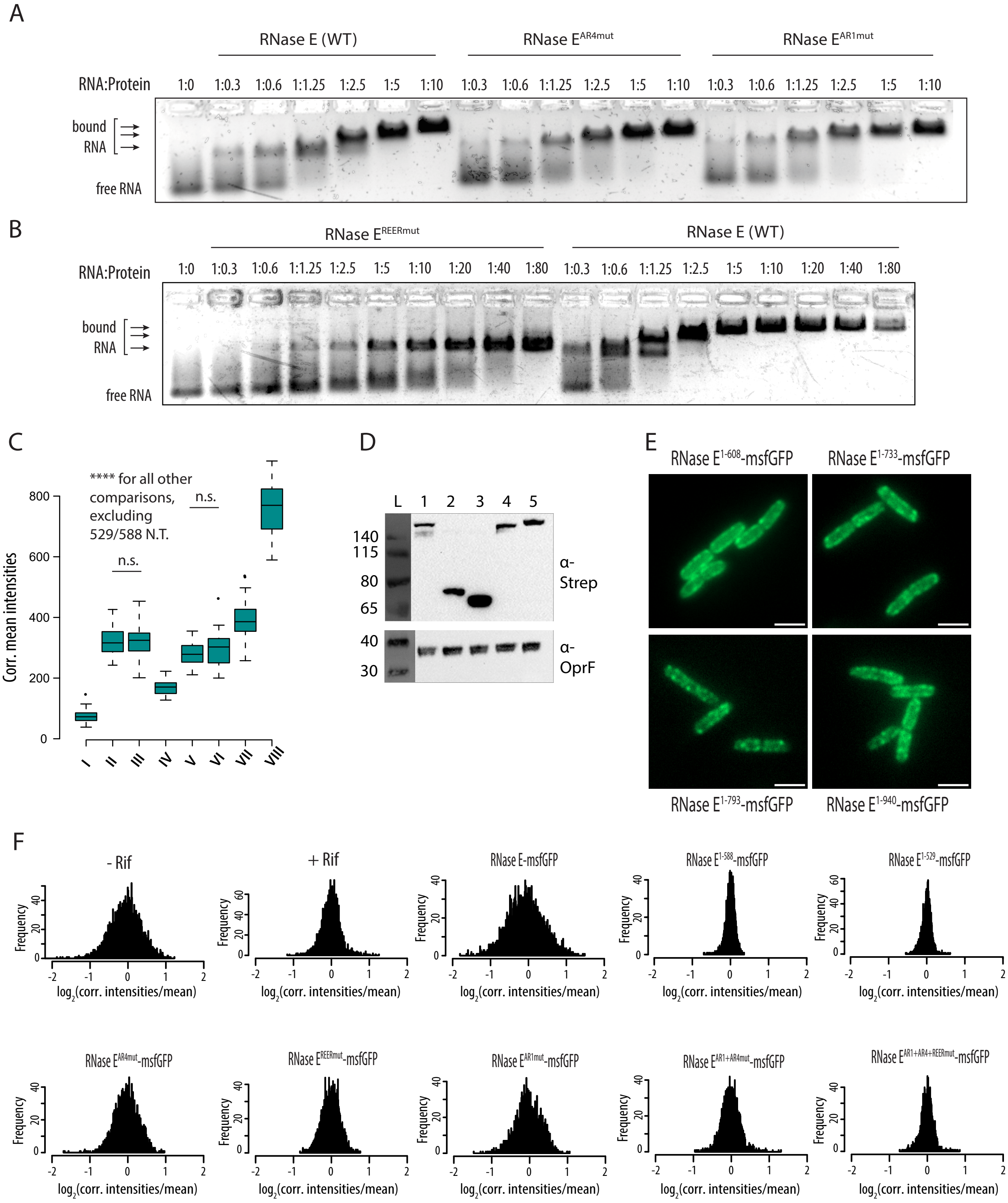

A
RNase E (WT)
RNase E$^{AR4mut}$
RNase E$^{AR1mut}$
RNA:Protein
1:0 1:0.3 1:0.6 1:1.25 1:2.5 1:5 1:10 1:0.3 1:0.6 1:1.25 1:2.5 1:5 1:10 1:0.3 1:0.6 1:1.25 1:2.5 1:5 1:10
bound RNA
free RNA
B
RNase E$^{REERmut}$
RNase E (WT)
RNA:Protein
1:0 1:0.3 1:0.6 1:1.25 1:2.5 1:5 1:10 1:20 1:40 1:80 1:0.3 1:0.6 1:1.25 1:2.5 1:5 1:10 1:20 1:40 1:80
bound RNA
free RNA
C
**** for all other comparisons, excluding 529/588 N.T.
n.s.
n.s.
Corr. mean intensities
0 200 400 600 800
I II III IV V VI VII VIII
D
L 1 2 3 4 5
140 115 80 65
40 30
α-Strep
α-OprF
E
RNase E$^{1-608}$-msfGFP
RNase E$^{1-733}$-msfGFP
RNase E$^{1-793}$-msfGFP
RNase E$^{1-940}$-msfGFP
F
- Rif
+ Rif
RNase E-msfGFP
RNase E$^{1-588}$-msfGFP
RNase E$^{1-529}$-msfGFP
RNase E$^{AR4mut}$-msfGFP
RNase E$^{REERmut}$-msfGFP
RNase E$^{AR1mut}$-msfGFP
RNase E$^{AR1+AR4mut}$-msfGFP
RNase E$^{AR1+AR4+REERmut}$-msfGFP
Frequency
log$_2$(corr. intensities/mean)

### Figure S5

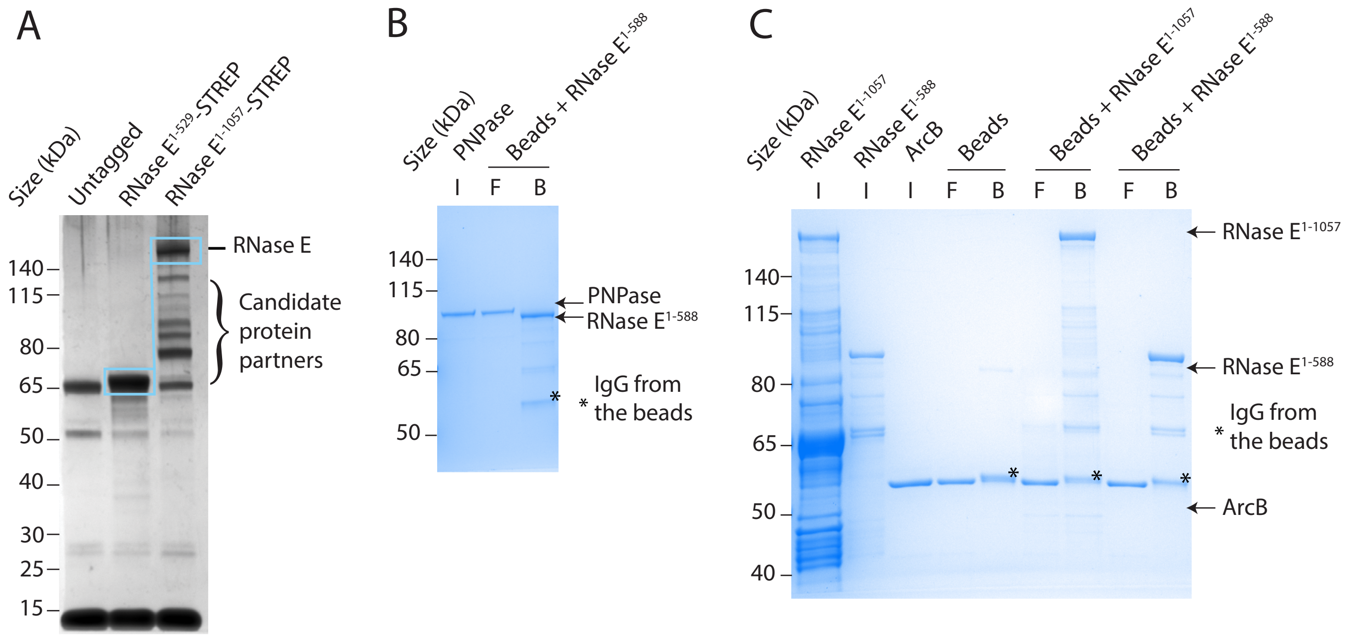

A
Size (kDa)
Untagged
RNase E1-529-STREP
RNase E1-1057-STREP
140
115
80
65
50
40
30
25
15
RNase E
Candidate protein partners
B
Size (kDa)
PNPase
Beads + RNase E1-588
I
F
B
140
115
80
65
50
PNPase
RNase E1-588
IgG from the beads
C
Size (kDa)
RNase E1-1057
RNase E1-588
ArcB
Beads
Beads + RNase E1-1057
Beads + RNase E1-588
I
I
I
F
B
F
B
F
B
140
115
80
65
50
40
RNase E1-1057
RNase E1-588
IgG from the beads
ArcB

### Figure S6

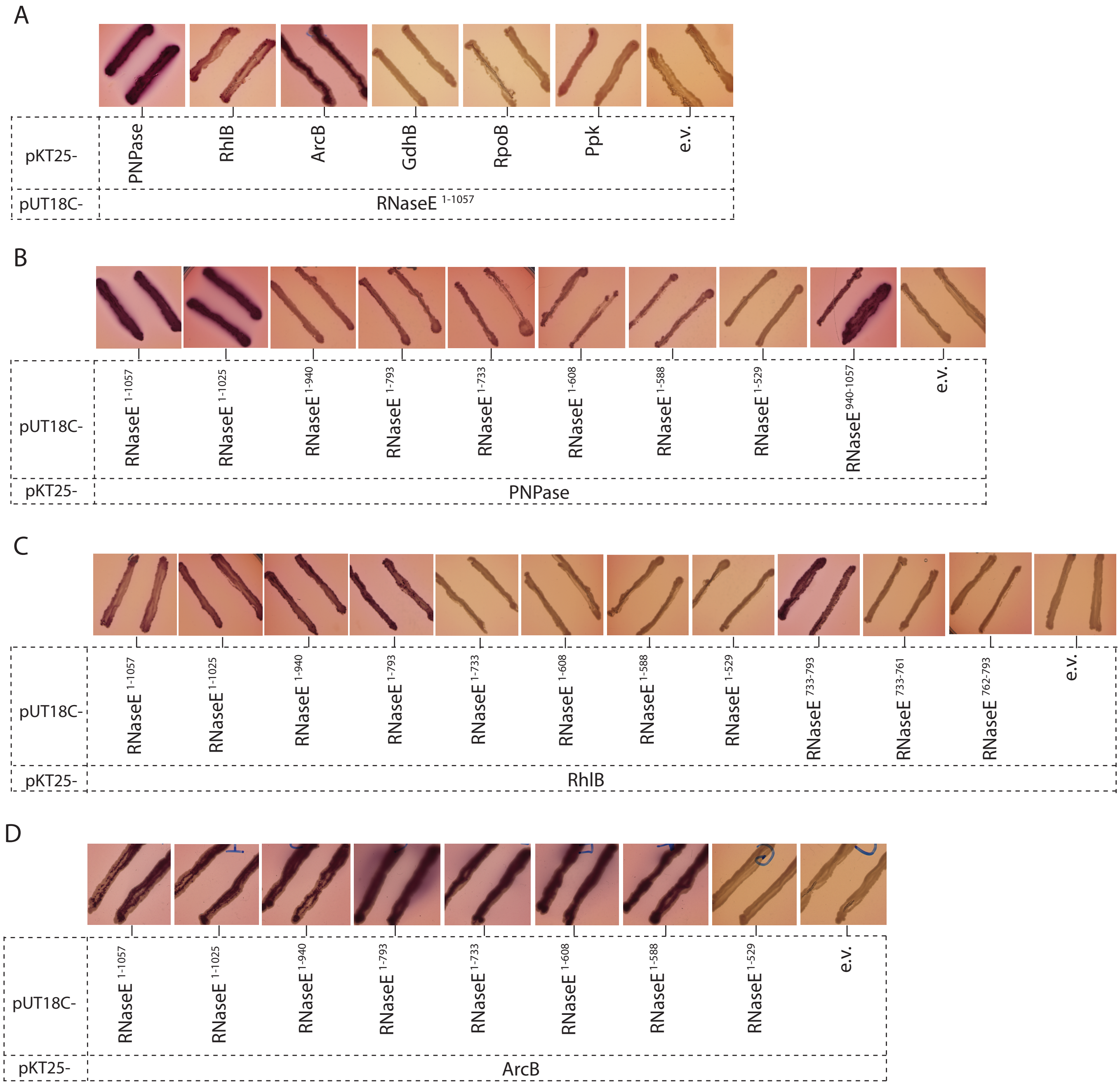
A
pKT25-
PNPase
RhlB
ArcB
GdhB
RpoB
Ppk
e.v.
pUT18C-
RNaseE 1-1057
B
pUT18C-
RNaseE 1-1057
RNaseE 1-1025
RNaseE 1-940
RNaseE 1-793
RNaseE 1-733
RNaseE 1-608
RNaseE 1-588
RNaseE 1-529
RNaseE 940-1057
e.v.
pKT25-
PNPase
C
pUT18C-
RNaseE 1-1057
RNaseE 1-1025
RNaseE 1-940
RNaseE 1-793
RNaseE 1-733
RNaseE 1-608
RNaseE 1-588
RNaseE 1-529
RNaseE 733-793
RNaseE 733-761
RNaseE 762-793
e.v.
pKT25-
RhlB
D
pUT18C-
RNaseE 1-1057
RNaseE 1-1025
RNaseE 1-940
RNaseE 1-793
RNaseE 1-733
RNaseE 1-608
RNaseE 1-588
RNaseE 1-529
e.v.
pKT25-
ArcB

### Figure S7

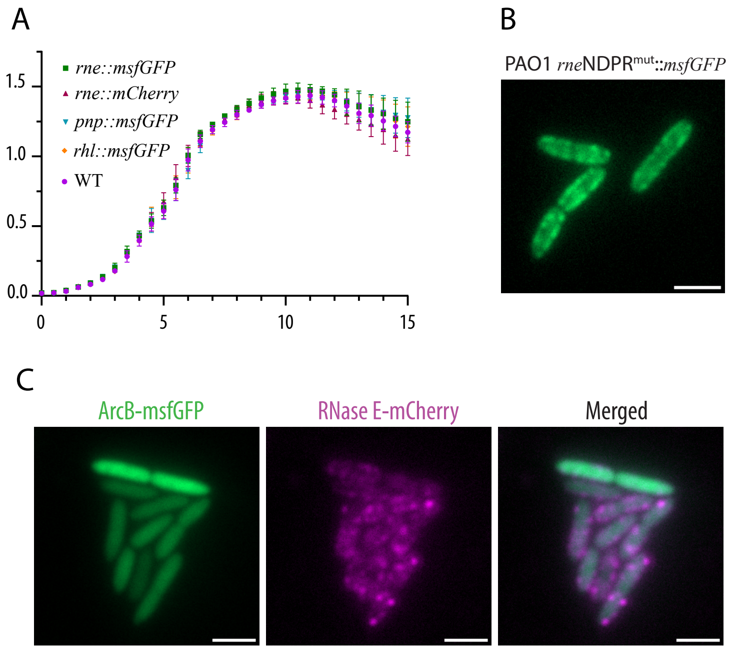
A
rne::msfGFP
rne::mCherry
pnp::msfGFP
rhl::msfGFP
WT
1.5
1.0
0.5
0.0
0
5
10
15
B
PAO1 rneNDPR$^{mut}$::msfGFP
C
ArcB-msfGFP
RNase E-mCherry
Merged

### Figure S8

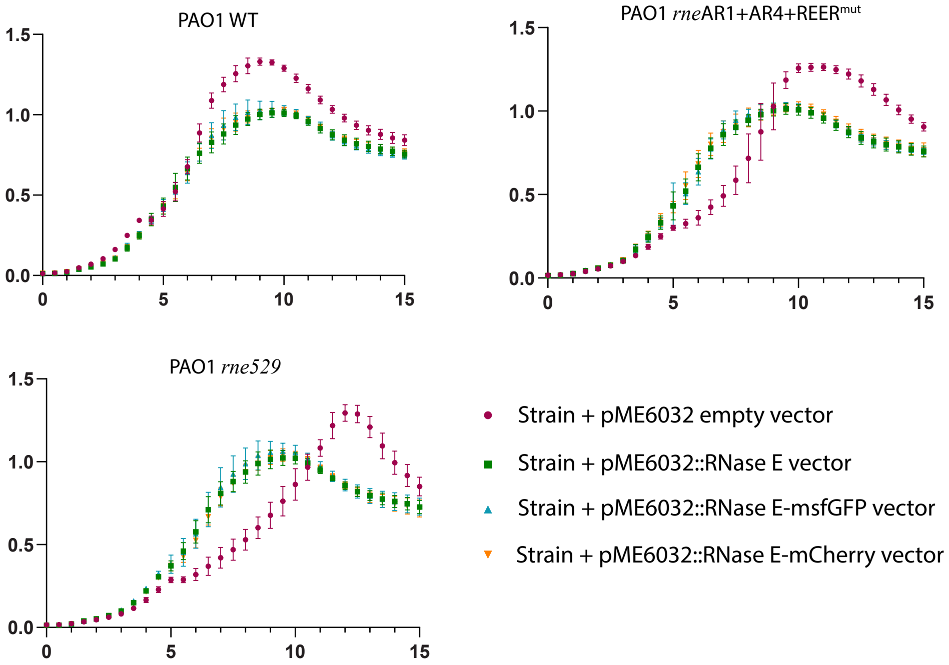
PAO1 WT
PAO1 rneAR1+AR4+REERmut
PAO1 rne529
1.5
1.0
0.5
0.0
0
5
10
15
Strain + pME6032 empty vector
Strain + pME6032::RNase E vector
Strain + pME6032::RNase E-msfGFP vector
Strain + pME6032::RNase E-mCherry vector

### Figure S9

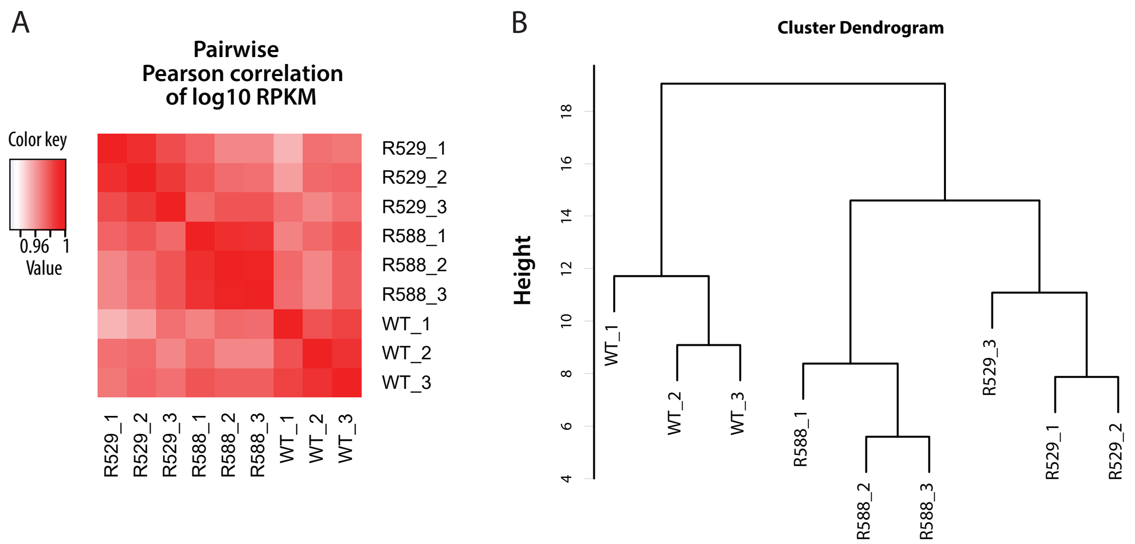
A
Pairwise
Pearson correlation
of log10 RPKM
Color key
0.96
1
Value
R529_1
R529_2
R529_3
R588_1
R588_2
R588_3
WT_1
WT_2
WT_3
B
Cluster Dendrogram
Height
4
6
8
10
12
14
16
18

Figure S10

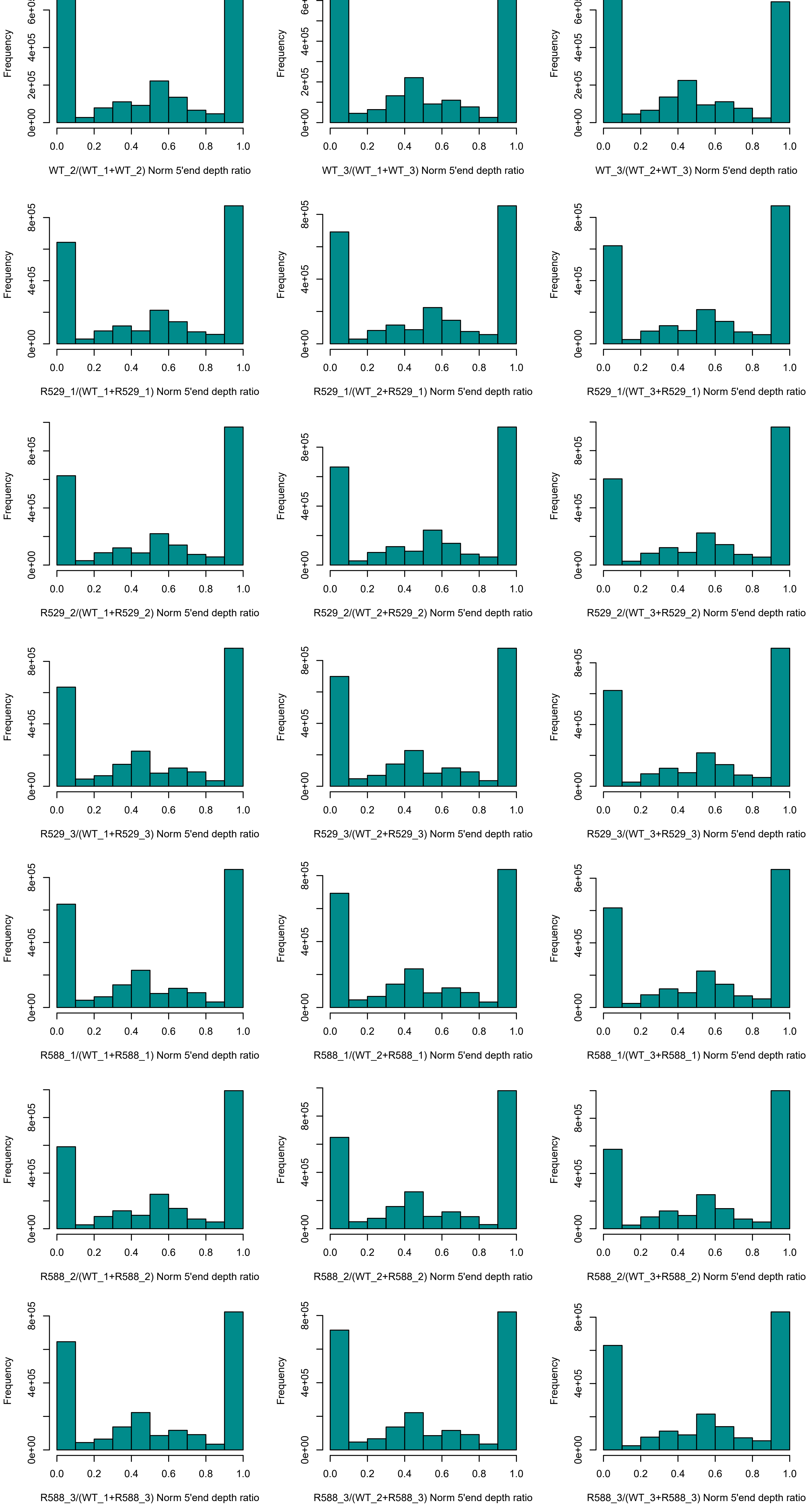

Frequency
0e+00
2e+05
4e+05
6e+05
8e+05
0.0
0.2
0.4
0.6
0.8
1.0
WT_2/(WT_1+WT_2) Norm 5'end depth ratio
WT_3/(WT_1+WT_3) Norm 5'end depth ratio
WT_3/(WT_2+WT_3) Norm 5'end depth ratio
R529_1/(WT_1+R529_1) Norm 5'end depth ratio
R529_1/(WT_2+R529_1) Norm 5'end depth ratio
R529_1/(WT_3+R529_1) Norm 5'end depth ratio
R529_2/(WT_1+R529_2) Norm 5'end depth ratio
R529_2/(WT_2+R529_2) Norm 5'end depth ratio
R529_2/(WT_3+R529_2) Norm 5'end depth ratio
R529_3/(WT_1+R529_3) Norm 5'end depth ratio
R529_3/(WT_2+R529_3) Norm 5'end depth ratio
R529_3/(WT_3+R529_3) Norm 5'end depth ratio
R588_1/(WT_1+R588_1) Norm 5'end depth ratio
R588_1/(WT_2+R588_1) Norm 5'end depth ratio
R588_1/(WT_3+R588_1) Norm 5'end depth ratio
R588_2/(WT_1+R588_2) Norm 5'end depth ratio
R588_2/(WT_2+R588_2) Norm 5'end depth ratio
R588_2/(WT_3+R588_2) Norm 5'end depth ratio
R588_3/(WT_1+R588_3) Norm 5'end depth ratio
R588_3/(WT_2+R588_3) Norm 5'end depth ratio
R588_3/(WT_3+R588_3) Norm 5'end depth ratio
